# Transgene directed induction of a stem cell-derived human embryo model

**DOI:** 10.1101/2023.06.15.545082

**Authors:** Bailey AT Weatherbee, Carlos W Gantner, Riza M Daza, Nobuhiko Hamazaki, Lisa K. Iwamoto-Stohl, Jay Shendure, Magdalena Zernicka-Goetz

## Abstract

The human embryo undergoes morphogenetic transformations following implantation into the uterus, yet our knowledge of this crucial stage is limited by the inability to observe the embryo *in vivo*. Stem cell-derived models of the embryo are important tools to interrogate developmental events and tissue-tissue crosstalk during these stages^1^. Here, we establish a human post-implantation embryo model comprised of embryonic and extraembryonic tissues. We combine two types of extraembryonic-like cells generated by transcription factor overexpression with wildtype embryonic stem cells and promote their self-organization into structures that mimic aspects of the post-implantation human embryo. These self-organized aggregates contain a pluripotent epiblast-like domain surrounded by hypoblast-and trophoblast-like tissues. We demonstrate that these inducible human embryoids robustly generate several cell types, including amnion, extraembryonic mesenchyme, and primordial germ cell-like cells in response to BMP signaling. This model also allowed us to identify an inhibitory role for SOX17 in the specification of anterior hypoblast-like cells^2^. Modulation of the subpopulations in the hypoblast-like compartment demonstrated that extraembryonic-like cells impact epiblast-like domain differentiation, highlighting functional tissue-tissue crosstalk. In conclusion, we present a modular, tractable, integrated^3^ model of the human embryo that will allow us to probe key questions of human post-implantation development, a critical window when significant numbers of pregnancies fail.

## Introduction

Human reproduction is remarkably inefficient, with an estimated 60% of pregnancies failing during the first two weeks following fertilization^4, 5^. Since the advent of *in vitro* fertilization, human embryos have been studied throughout the first week of development^1, 6^. However, the second week, which includes implantation into the uterus and preparation for gastrulation, remains a ‘black box^5^’. The human blastocyst at 5-6 days post-fertilization consists of the outermost trophectoderm, precursor of the placenta, and inner cell mass, which gives rise to both the embryonic epiblast and the yolk sac precursor, the hypoblast. Between 7-8 days post-fertilization the blastocyst implants into the endometrium and the epiblast polarizes and transitions from the naïve state of pluripotency to the primed state^1^. A central amniotic cavity forms within the epiblast, separating the dorsal amniotic epithelium and the ventral epiblast, which maintains pluripotency and gives rise to the embryo proper^1^. The trophectoderm develops into several trophoblast subtypes following implantation^7^ and the hypoblast forms the primary, and then secondary, yolk sac. A subset of cells in the hypoblast maintains expression of NODAL, BMP and WNT inhibitors, safeguarding the future anterior epiblast from posteriorizing signals during primitive streak formation, marked by upregulation of BRY/TBXT^2^. An additional extraembryonic tissue, the extraembryonic mesenchyme, is located between the inner cell mass-derived tissues and the trophoblast, however, the origin of these cells remains unclear^8^.

Recent work in mouse embryos established conditions amenable to human embryo *in vitro* culture through implantation, opening this developmental black box for the first time^9–12^. We and other groups have used this system to characterize major developmental events, including formation of the anterior hypoblast domain, specification of trophoblast subtypes and transitions in epiblast pluripotency state^2, 7,13–15^. However, mechanistic work in the human embryo remains challenging. Thus, stem cell-derived models of the human embryo will serve as an important and complementary tool to understanding this crucial period of our development. Several groups have reported the generation of blastocyst-like structures derived from human embryonic stem cells (hESCs)^16–20^. These ‘blastoids’ resemble the pre-implantation embryo but develop poorly to post-implantation stages. Other models, including gastruloids, 2D micropatterns and embryoid bodies, are capable of modelling aspects of post-implantation development^21–24^. However, these models are derived entirely from hESC, lack extraembryonic tissues, and do not recapitulate embryo morphology. A recent study, which combines epiblast-like spheroids with BMP4-treated hESCs expressing a mixture of extraembryonic markers, marks a step towards integrated models of the post-implantation embryo^25^. However, this model does not exhibit self-organization of an epiblast-like compartment in the context of extra-embryonic tissues until after lumenogenesis, and the BMP4 treated hESCs are not correlated to targeted extraembryonic lineages.

Several protocols have been developed to derive trophoblast and hypoblast cells from hESCs^26–31^. Importantly, the pluripotent state influences the developmental trajectory of differentiated cells^26,27^. The derivation of lineage-specific cell lines offers scope for modelling these tissues *in vitro*. However, generating a modular, integrated model system that includes both embryonic and extraembryonic tissues has proven challenging. This may be due to the opposing signaling pathway modulators required for in hESC culture, hypoblast-like cell differentiation, and trophoblast-like cell differentiation. Moreover, while tissue-tissue crosstalk is an advantage of integrated model systems, generation of embryoids in media containing exogenous factors may compromise tissue-driven self-organization. Given these limitations, and the ability of tri-lineage models to mimic mouse development^32–34^, we utilized the approach of overexpressing transcription factors that can drive generation of extraembryonic-like cells without the need for exogenous factors.

We show that aggregates of induced extraembryonic-like lineages and wildtype hESC are capable of self-organisation into embryo-like structures, which mimic several hallmarks of post-implantation development, including lumenogenesis, amniogenesis, primordial germ cell formation, and specification of the anterior hypoblast. Importantly, these inducible human embryoids are modular, do not rely on exogenous signaling factors, and are amenable to genetic perturbation.

## Results

### Transcription factor-mediated induction of extraembryonic lineages

Mouse ESCs can be induced to generate extraembryonic endoderm or trophoblast through expression of Gata4 or Cdx2, respectively, and these cells are capable of modelling development in a 3D embryoid models^33–35^. Thus, we first identified factors that can similarly upregulate extraembryonic gene programs in human ESC. Notably, pluripotent state, in part, dictates differentiation potential from ESCs^26,27,29,30,36^. Therefore, we overexpressed candidate transcription factors in hESCs across the pluripotency spectrum to assess which factors are able to program hESCs to trophoblast or hypoblast-like cells. We first integrated published single cell RNA-sequencing data of human embryos cultured until gastrulation^2,13, 37–40^ and used the computational tool SCENIC to score the predicted activity of transcription factors enriched in the epiblast, trophoblast or hypoblast to generate a putative gene regulatory network (**Extended Figure 1A**)^41^. As expected, we found that *SOX2*, *NANOG* and *POU5F1* (OCT4) showed high predicted activity in the epiblast. Transcription factors including *GATA4*, *GATA6*, *SOX17* and *FOXA2* were particularly active in the hypoblast and *GATA3*, *NR2F2*, *GATA2* and *TFAP2C* (also known as AP2γ) showed enriched activity in the trophoblast (**Extended Figures 1A-B**). The overexpression of *GATA6* or *SOX17* has been shown to drive endodermal gene programs from primed hESCs^42, 43^, therefore, we selected them as candidates to program hESCs to become hypoblast-like. Similarly, *GATA3* and *TFAP2C* have been reported to share high chromatin co-occupancy during differentiation of hESCs into trophoblast stem cells^44^. This, together with their high predicted activity in trophoblast, led us to select *GATA3* and *TFAP2C* as candidates to drive hESCs to become trophoblast-like. We generated hESCs with doxycycline-inducible individual or combined transgenes for the transcription factors of interest: namely *GATA6* and *SOX17* to program hESCs to hypoblast-like cells (**Figure 1A**); and *GATA3* (together with an *eGFP* also driven by doxycycline) and *TFAP2C* to program hESCs to trophoblast-like cells (**Figure 1B**). Transcription factor induction was validated following doxycycline administration for 24 hours (**Figures 1A-B**).

**Figure 1.**
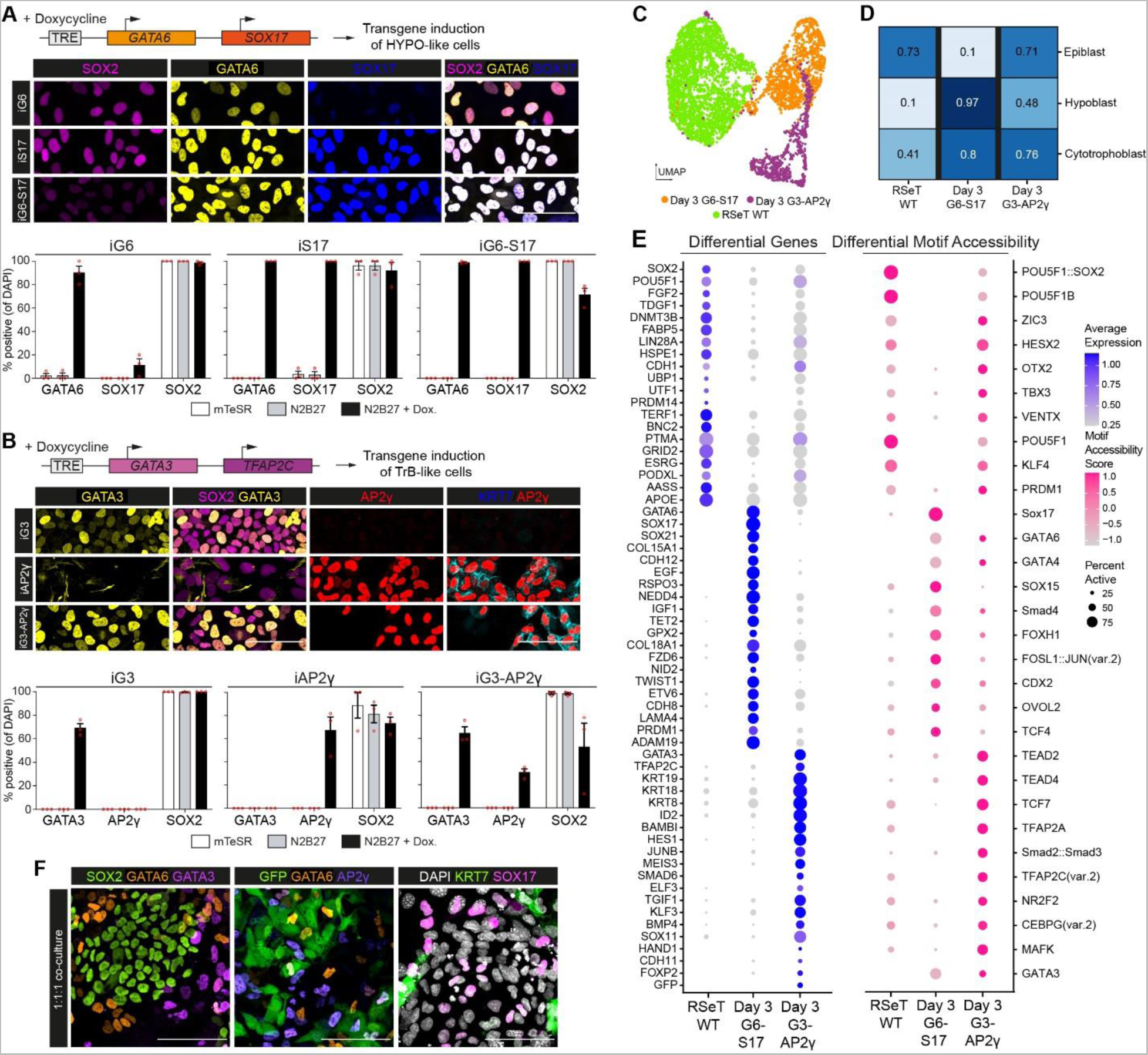
Validation of extraembryonic-like induction. (A) Generation of inducible GATA6 (iG6) and/or SOX17 (iS17) hESCs and validation after 24 hours of doxycycline addition in basal N2B27. N=3 independent exerpiments with 551 iG6, 550 iS17, 707 G6-S17 cells (B) Generation of inducible GATA3 (iG3) and/or AP2γ (iAP2γ) hESCs and validation after 24 hours of doxycycline addition in basal N2B27. N=3 independent experiments with 1456 iG3, 409 iAP2γ, 782 iG3-AP2γ cells (C) Visualization of Uniform Manifold Approximation and Projection (UMAP)-based dimensional reduction of sequenced wildtype (RSeT WT), inducible GATA6-SOX17 (Day 3 iG6-S17) and inducible GATA3-AP2γ (Day 3 iG3-AP2γ) RSeT hESCs after 3 days of doxycycline induction. (D) Logistic regression analysis and comparison of cells to human post-implantation embryo populations. Cell line data projected onto human embryo data from Molè et al., 2021. (E) Selected differentially expressed genes from RNA-sequencing (blue; left) and predicted differential motif accessibility from ATAC-sequencing scored by chromVAR (pink; right) for wildtype, inducible GATA6-SOX17 and inducible GATA3-AP2γ RSeT hESCs after 3 days of doxycycline induction. (F) Validation of wildtype, inducible GATA6-SOX17 and inducible GATA3-AP2γ RSeT hESC co-culture in 2D.

To assess the capacity of the selected candidate transcription factors to program hESCs to extra-embryonic-like cell identity, we overexpressed them – singly and in combination – in cells across the naïve-to-primed pluripotency spectrum. We cultured cells using three established starting conditions: PXGL, which supports pre-implantation-like naïve hESCs; RSeT, which generates intermediate peri-implantation-like pluripotent hESCs; and conventional mTeSR1 conditions to maintain post-implantation-like primed hESCs. After three days of transcription factor overexpression in basal N2B27 media, we observed significant differences in extraembryonic gene induction using both individual or combined transgenes and starting from different pluripotency states, at both the protein and mRNA level (**Extended Figure 1C-D, Extended Figure 2**). In hypoblast-like cell induction, GATA6 overexpression did not drive SOX17 expression after 3 days of induction in N2B27 from RSeT or PXGL conditions, but SOX17 overexpression resulted in robust GATA6 upregulation across starting pluripotency state conditions (**Extended Figure 1C, 2A-B**). FOXA2 expression was consistently upregulated after combined GATA6 and SOX17 induction from primed and RSeT, but not from PXGL conditions (**Extended Figure 2A-B**). These data indicate that while GATA6 and SOX17 can indeed drive endodermal gene programs, the regulation of specific downstream targets differs depending on the initial pluripotency state.

The AP2γ transgene appeared particularly effective in upregulating GATA2 and CK7 expression when driving trophoblast-like cell formation. However, induction of AP2γ alone resulted in cell death and loss of transgene expression in primed, but not RSeT or PXGL, cells (**Extended Figure 2C-D**). Combined induction of GATA6 and SOX17 or GATA3 and AP2γ resulted in consistent downregulation of pluripotency markers, including NANOG, SOX2 and OCT4 (**Extended Figure 1C-D, Extended Figure 2A-D)**.

We hypothesized that RSeT hESCs are the best starting cell type to generate our stem cell-derived human post-implantation embryo model because they: (1) represent a peri-implantation stage of development^27^; (2) express low levels of amnion-specific genes during trophoblast-like cell induction compared to primed cells (**Extended Figure 1D, Extended Figure 2C-D**); and (3) are known to more readily differentiate to peri- and post-implantation yolk sac-like endoderm cells as compared to PXGL cells^26, 27^. For these reasons, and the synergistic action of dual induction of candidate transcription factors, we used RSeT hESCs with inducible GATA6 and SOX17 transgenes for hypoblast-like cell induction and used cells with inducible GATA3 and AP2γ transgenes for trophoblast-like cell induction in subsequent experiments. We compared transcription factor overexpression-based induction of extraembryonic gene programs to established exogenous factor-based hypoblast-like cell and trophoblast-like cell directed differentiation protocols. Dual induction of GATA6 and SOX17 from RSeT cells in basal media induced endodermal gene expression equivalent to directed differentiation protocols in yolk sac-like cell differentiation conditions (Activin-A, CHIR99021, and LIF) (**Extended Figures 3A-B**). Dual induction of GATA3 and AP2γ from RSeT cells in basal media induced trophoblast gene expression, albeit at varying levels when compared to directed trophoblast differentiation protocols using MEK/ERK inhibition (PD0325901) and ALK5/7 inhibition (A83-01) with or without Hippo signaling pathway inhibition (lysophosphatidic acid; LPA) (**Extended Figures 3C-D**).

To further characterize RSeT hESCs induced to express GATA6-SOX17 or GATA3-AP2γ, we carried out single-cell 10x multiome sequencing and assessed transcriptomic and chromatin accessibility simultaneously. When this single-cell data was visualized using uniform manifold approximation and projection (UMAP), cells clustered based on cell type (**Figure 1C**), and the application of a logistic regression framework showed that the wildtype, inducible GATA6-SOX17, and inducible GATA3-AP2γ RSeT hESCs had the highest similarities to the epiblast, hypoblast, and cytotrophoblast of the post-implantation embryo, respectively (**Figure 1D**). Analysis of differentially expressed genes and differentially accessible motifs scored by chromVAR^45^ revealed similar embryonic and extraembryonic dynamics (**Figure 1E, Extended Figure 4, Extended Table 1**). Specifically, we detected enriched expression and motif accessibility scores of pluripotency and epiblast markers in RSeT hESCs (*SOX2*, *POU5F1*, *TDGF1*); of hypoblast markers in GATA6-SOX17 inducible cells (*RSPO3*, *IGF1*, *PRDM1*); and of trophoblast markers (*KRT19, KRT18, BMP4*), including Hippo, WNT and TGFβ pathway components in the GATA3-AP2γ inducible cells. Together, these data demonstrate that transcription factor-mediated induction of extraembryonic cell fate from RSeT hESCs readily drives hypoblast- or trophoblast-like gene programs without the need for exogenous factors. This allows us to overcome the challenge presented for successful coculture of these cell types by their conflicting culture media requirements. Indeed, 1:1:1 co-culture of wildtype RSeT hESCs with inducible GATA6-SOX17 and inducible GATA3-AP2γ RSeT hESCs demonstrated good survival and mixed identity after transgene induction for 3 days **(Figure 1F**).

### Self-assembly of a tri-lineage 3D post-implantation model

As all three RSeT hESC-derived cell types – wildtype, GATA6-SOX17 inducible and GATA3-AP2γ inducible – can be cocultured in N2B27 medium, we induced expression of the selected transcription factors with doxycycline for 3 days to generate hypoblast- and trophoblast-like cells and subsequently aggregated cell mixtures in Aggrewell dishes (**Figure 2A**).

**Figure 2.**
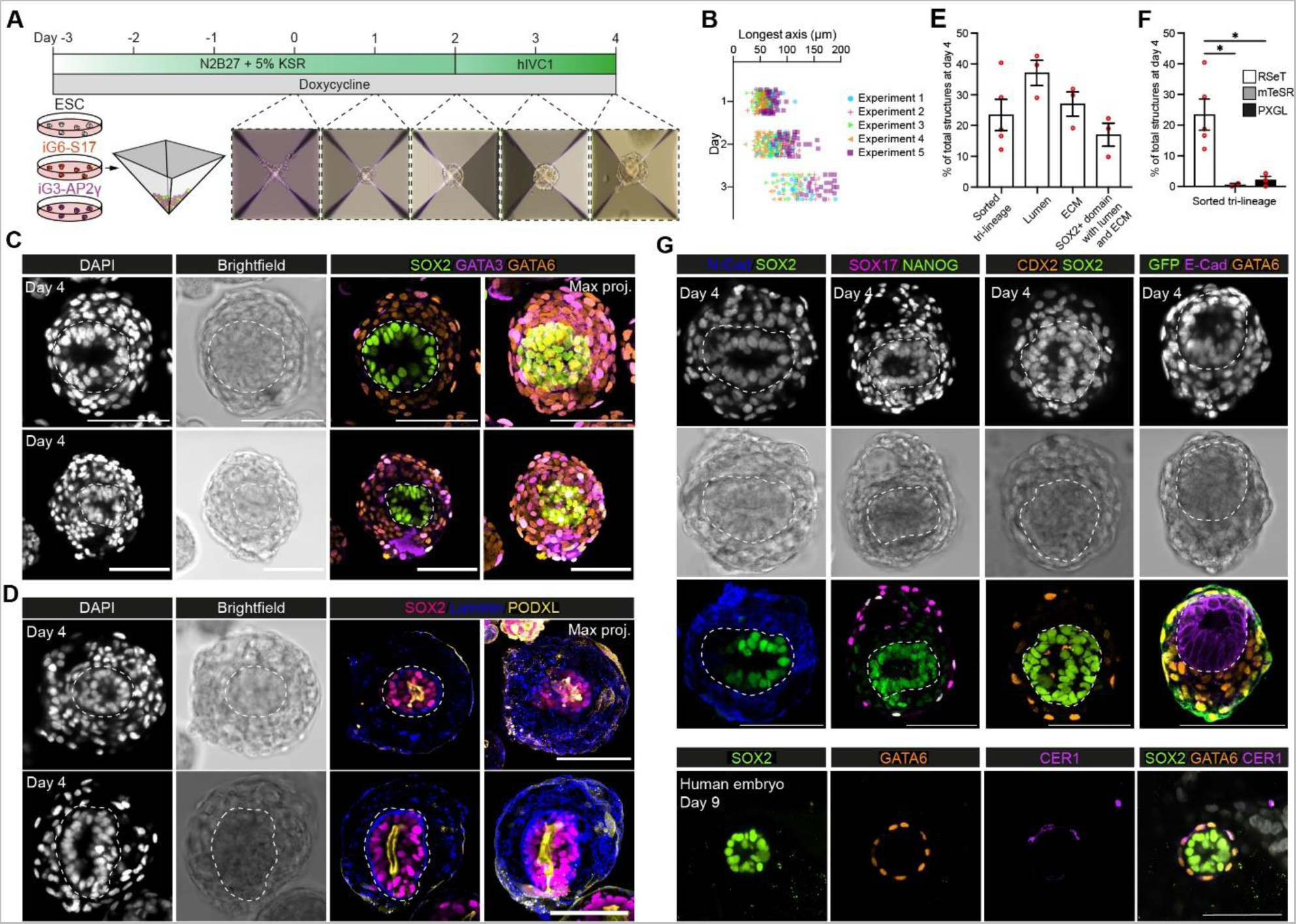
Generation of inducible post-implantation human embryoids. (A) Overview of experimental protocol to generate inducible human embryoids by combining wildtype RSeT hESC with inducible extraembryonic-like cells. To generate hypoblast-like cells, inducible GATA6-SOX17 (iG6-S17) was used. To generated trophoblast-like cells, inducible GATA3-AP2γ (iG3-AP2γ) was used. Extraembryonic-like cells were induced for 3 days before aggregation at Day 0. (B) Size of cell aggregates between day 1 to day 3 following aggregation. N=5 independent experiments with 175 day 1, 171 day 2, and 91 day 3 structures. (C) At 96 hours after aggregation, structures demonstrate clear self-organization with an inner SOX2-positive domain surrounded by sequential layers of GATA6-positive and GATA3-positive extraembryonic-like populations. (D) Inducible human embryoids demonstrate clear apicobasal polarity within the inner SOX2-positive domain, including PODXL-positive lumen formation and basal deposition of laminin between the SOX2-positive epiblast-like and hypoblast-like compartments. (E) Quantification of inducible human embryoid formation efficiency and organization. N=5 indpedent experiments with 952 structures for tri-lineage, 3 independent experiments with 506 structures for lumen and ECM efficiency. (F) Quantification of inducible human embryoid formation across starting pluripotency states. N=5 independent experiments with 1189 structures quantified. (G) The hypoblast-like domain expresses N-Cadherin and SOX17. Cells derived from inducible GATA3-AP2γ (which expresses GFP) show clear outer localization. (H) Representative image of an in vitro cultured human embryo at 9 days post-fertilization showing clear lumenized SOX2 domain surrounded by a layer of GATA6-postive cells. A subset of GATA6-positive cells expresses the anterior hypoblast marker CER1. Scale bars = 100mm. Statistics: (F) one-way ANOVA with Dunnett’s multiple comparisons test. *p<0.05.

We plated a ratio of 1:1:2 wildtype:GATA6-SOX17-inducible:GATA3-AP2γ-inducible cells with 3.6 x 10^4^ cells per Aggrewell-400 24-well in N2B27 media (approximately 8:8:16 cells per microwell). Cells aggregated within 24 hours, and by 48 hours post-aggregation we observed clear distinctions between inner and outer cellular domains by brightfield (**Figure 2A**). At 48 hours post-aggregation, the media was changed to post-implantation human embryo media (hIVC1)^2, 12, 13^. Incubation with doxycycline continued throughout the whole period of culture (**Figure 2A**). Cells aggregated and proliferated consistently across experiments (**Figure 2B**). Four days post-aggregation, the cell aggregates had self-organized into structures with a SOX2-positive, epiblast-like domain containing a central lumen; an outer single layer of GATA3-positive putative trophoblast-like cells; and an intermediate putative hypoblast-like domain of GATA6-positive cells between inner lumenized domain and outer layer (**Figure 2C**). The inner SOX2-positive domain exhibited apicobasal polarity with apical expression of the luminal marker PODXL, and basal deposition of laminin (**Figure 2D**). Similar to experiments in which mouse embryonic and extraembryonic stem cells were aggregated to generate a post-implantation mouse embryo model, these structures did not transit through a blastocyst-like morphology prior to forming post-implantation-like structures^32–34, 46–48^. The efficiency of inducible human embryoid formation (defined as aggregates containing an organized SOX2-positive domain, surrounded by concentric layers of GATA6-positive and GATA3-positive cells) was approximately 23% (**Figure 2E**). By contrast, when using primed mTeSR1 or naïve PXGL hESCs as the starting pluripotency state for constitutive wildtype, GATA6-SOX17 inducible, and GATA3-AP2γ inducible cells, the efficiency of organized, multi-lineage structure formation was less than 5% (**Figure 2F**).

Our inducible human embryoids expressed several other lineage markers in an organized manner, including N-Cadherin and SOX17 in the putative hypoblast-like compartment and CDX2 in the outer layer of putative trophoblast-like cells (**Figure 2G**). We also observed structures with SOX17 and/or GATA6 expression within outer GATA3-AP2γ-induced cells (marked by eGFP), which may reflect the reported tendency of peripheral cells to adopt endodermal identities in embryoid bodies^49, 50^. The epiblast-like inner compartment expressed SOX2, NANOG and E-Cadherin and maintained pluripotent and epithelial identity akin to the human embryo (**Figure 2G**). These data demonstrate that RSeT hESC-derived epiblast-, hypoblast- and trophoblast-like cells are able to self-organize into embryo-like structures reminiscent of the post-implantation human embryo at 8-9 days post-fertilization (**Figure 2H**).

### Generation of amnion, primordial germ cells, and extra-embryonic mesenchyme

To gain insight into whether our human embryo model develops gene expression and chromatin accessibility patterns that align to the natural human embryo, we performed single-cell multiome RNA and ATAC sequencing on human embryo models at 4, 6, and 8 days post-aggregation (**Figure 3A**). We selected individual structures for sequencing based on their development of the three tissues: (1) an inner, epithelial domain; (2) an intermediate domain surrounding the central epithelium; and (3) an outer GFP-positive cell layer (**Extended Figure 5A**). We used these criteria because these structures represented self-organized stem cell-derived aggregates that resembled aspects of the post-implantation human embryo. After quality control to filter out cell barcodes with low mapped reads to either RNA or chromatin, 5217 cells were included in the analysis spanning all three timepoints (**Figure 3B**). To assign clusters without bias, we used scmap^51^ to project our dataset onto human^2, 52^ and cynomolgus macaque^53–55^ datasets spanning peri-implantation to gastrula stages (**Figure 3B and Extended Figure 5B**). This analysis allowed us to project gene expression signatures from previously annotated cynomolgus macaque cell type clusters including epiblast-, amnion-, visceral endoderm-, extraembryonic mesenchyme- and gastrulation-like populations onto our human embryo model. The recently published multiome-based velocity inference program multivelo^56^ was used to predict lineage relationships from combined RNA and chromatin velocity measurements. We found that the inferred differentiation time correlated well with the transition of structures from day 4 to day 8 post-aggregation. These data, in combination with canonical marker expression, allowed us to annotate the cell types in our human embryo model (**Figure 3B and Extended Figure 5C**). We identified clusters resembling embryonic late-epiblast (L-EPI: *TDGF1, SOX2, NANOG-*positive), amnion (AM-1: *TFAP2A, ID1-*positive; AM-2: *ISL1, TFAP2C*-positive; and AM-3: *GABRP, VTCN1, GRHL1, MEIS1*-positive), mesoderm (MESO-1: *TBXT+, MESP1*-positive; MESO-2: *MIXL1, CER1, SNAI1, EOMES*-positive), extraembryonic mesenchyme (EXMC: *POSTN, COL6A3, IGF2, TBX20-* positive) and hypoblast/visceral endoderm (HYPO/VE: *BMP6, CDH2, HNF1B, FOXA2*-positive) (**Figure 3C and Extended Figure 5C-D**). We did not identify a distinct trophoblast-like cluster derived from the GFP-positive inducible GATA3-AP2γ cells, despite their presence as an outer layer within the inducible human embryoids (**Figure 3A, Extended Figure 5E**). Given the aberrant upregulation of endodermal markers after aggregation, inducible GATA3-AP2γ-derived cells were not likely to represent *bona fide* trophoblast. Nevertheless, these inducible human embryoids generated several cell types which do not appear to be formed when human embryos are cultured *in vitro* through implantation, including amnion and extraembryonic mesenchyme. This is in agreement with immunofluorescence analysis that demonstrated that the inner, SOX2-positive domain exited pluripotency by day 6 post-aggregation and upregulated amnion markers, including CDX2 and ISL1. In addition, by day 8 post-aggregation, the inner domain upregulated mature amnion markers VTCN1 and HAND1 (**Figure 3D, Extended Figure 6A-B**). The GATA6-positive domain also co-expressed HAND1, supporting the presence of extraembryonic mesenchyme (**Extended Figure 6A-B**). A subset of GATA6-positive cells showed high co-expression of FOXF1 or TBX20, highlighting the two populations within the intermediate domain of our human embryo model: hypoblast cells (GATA6-positive, FOXF1 or TBX20-low) and extraembryonic mesenchyme (GATA6-positive, FOXF1 or TBX20-high) (**Extended Figure 6C-F**).

**Figure 3.**
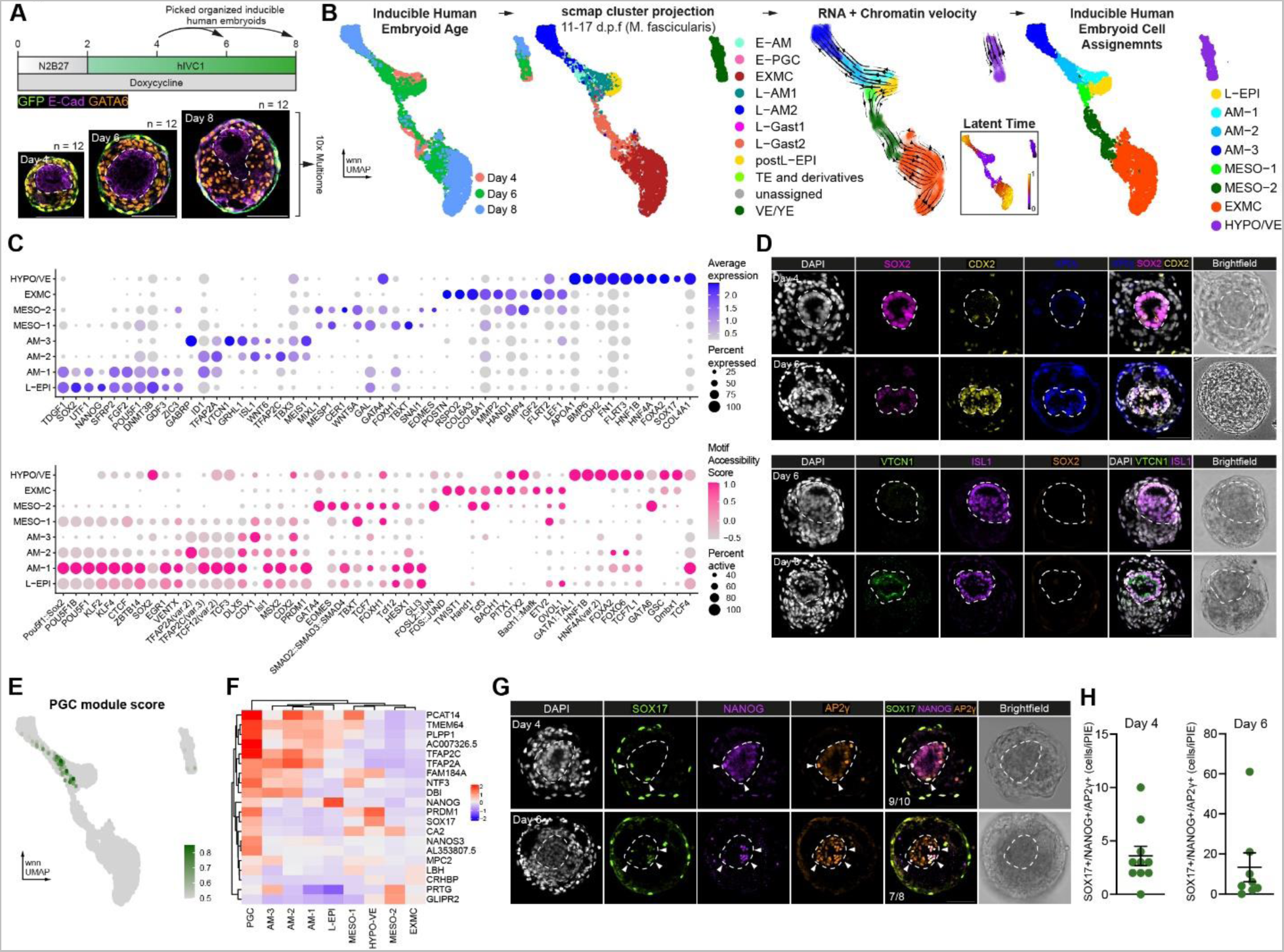
Differentiation of extraembryonic mesenchyme, amnion, and primordial germ cells. (A) Schematic of extended culture protocol of inducible human embryoids and sampling for combined single-cell RNA and single-cell ATAC sequencing using the 10x platform. (B) Cells were annotated based on transcriptional projection to multiple human and non-human primate embryo datasets using scmap in conjunction with RNA and chromatin velocity. (C) Selected differentially expressed genes in the RNA-sequencing data (top, blue) and predicted differentially accessible motifs scored by chromVAR on the ATAC-sequencing data (bottom, pink) across clusters. (D) Inducible human embryoids downregulated SOX2 and upregulated CDX2, ISL1 at day 6 and VTCN1 at day 8, indicative of robust amnion differentiation and maturation. (E) Module scoring for primordial germ cell marker genes (Jo et al., 2021). (F) Heatmap of selected primordial germ cell genes across clusters. (G) Immunofluorescence identification of SOX17/NANOG/AP2γ triple-positive primordial germ cell-like cells in inducible human embryoids highlighted by arrowheads. (H) Quantification of SOX17/NANOG/AP2γ triple-positive (+) cells at days 4 (n=10 embryoids) and 6 (n=10 embryoids). N=2 independent experiments. Scale bars = 100µm.

Recent reports have postulated that both amnion and primordial germ cells, the precursors to gametes, are at least partially generated from a bipotent progenitor *in vitro*^57–59^. We therefore asked whether such progenitors or their progeny were specified in our human embryo model. We first assigned a primordial germ cell module score based on expression of genes identified in human primordial germ cell-like cells differentiated *in vitro*^60^. We could identify cells with transcriptomes resembling primordial germ cell-like cells mostly throughout the amnion clusters AM-1, AM-2, and a small number within the mesodermal cluster MESO-1 **Figure 3E**). Putative primordial germ cell-like cells in inducible human embryoids expressed *TFAP2A* (also known as AP2α), a crucial marker of bipotent amnion and primordial germ cell-like cell progenitors^57–59^. In contrast to other cells in AM-1 and AM-2 clusters, primordial germ cell-like cells expressed the pluripotency marker *NANOG* and primordial germ cell markers *PRDM1* (also known as BLIMP1) and *NANOS3* (**Figure 3F**). Immunofluorescence analysis of a canonical set of human primordial germ cell markers^57^ confirmed that SOX17/NANOG/AP2γ triple-positive primordial germ cell-like cells were observed by day 4 post-aggregation and increased in number by day 6 post-aggregation (**Figures 3G-H**). These data demonstrate that our human embryo model specifies *bona fide* germ cells. This primordial germ cell-like cell specification is concomitant with amnion-like cell formation and occurs within the inner epiblast-like compartment, supporting the existence of a bipotent progenitor for these two lineages.

### The epiblast-like domain differentiates to amnion and extraembryonic mesenchyme in response to BMP signaling

Both amnion and extraembryonic mesenchyme are thought to arise in response to BMP signaling in the primate embryo^8, 53, 61, 62^. To understand if this is consistent in our human embryo model, we examined the expression of downstream BMP response genes *ID1-4*^63^. *ID1* and *ID4* were upregulated during amnion formation, while *ID3* and *ID2* were enriched in both amnion and extraembryonic mesenchyme trajectories, indicating that BMP signaling was likely active (**Figure 4A**)^63^. In support of this, SMAD5 motif accessibility based on the ATAC-sequencing (scored by chromVAR^45^) was high in both trajectories while SMAD2::SMAD3::SMAD4 motif accessibility score, a downstream target of Activin-NODAL signaling, was not (**Figure 4B**). In line with this observation, a high BMP and low NODAL signaling environment has been implicated recently in amnion differentiation of marmoset ESCs^64^ and in hESC to extraembryonic mesenchyme differentiation^62^, suggesting that similar dynamics may drive differentiation of these populations within inducible human embryoids.

**Figure 4.**
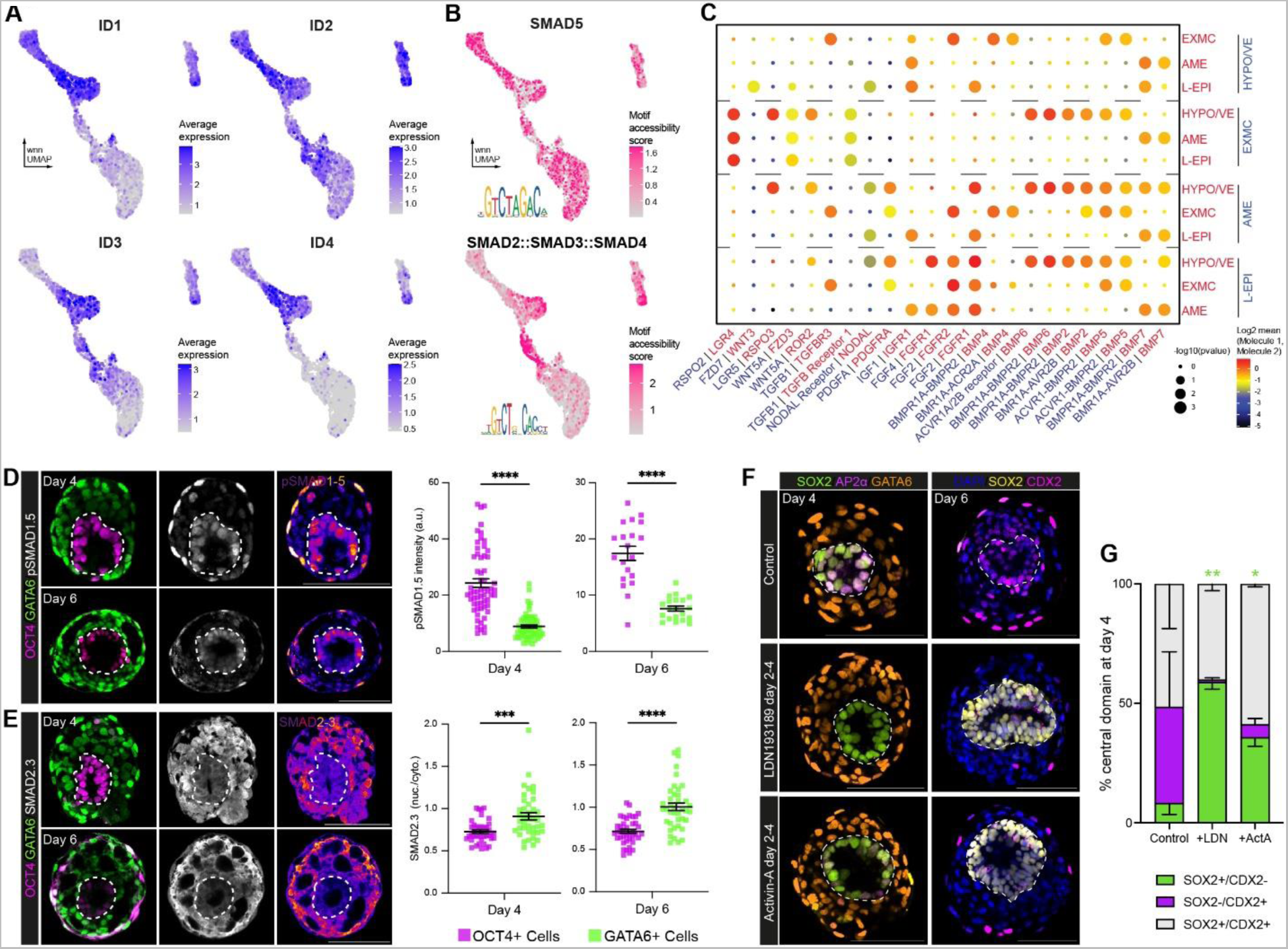
BMP signaling drives amnion specification in inducible human embryoids. (A) Expression of *ID1*, *ID2*, *ID3* and *ID4* transcripts, downstream targets of BMP signaling, in sequencing of post-implantation embryo models. (B) chromVAR-based motif accessibility scores of SMAD5 and SMAD2::SMAD3::SMAD4, effectors of BMP and NODAL signaling, respectively. (C) Predicted ligand-receptor pairings in inducible human embryoids generated by CellPhoneDB (Efremova et al., 2020). (D) Representative immunofluorescence staining and quantification of representative inducible human embryoids at days 4 (n=120 cells) and 6 (n=80 cells) showing heightened pSMAD1.5 in the epiblast-like domain. N=3 independent experiments (E) Representative immunofluorescence staining and quantification of SMAD2.3 in representative inducible human embryoids at days 4 (n=80 cells) and 6 (n=80 cells). N=2 independent experiments. (F) Inhibition of BMP signaling blocks exit from pluripotency and upregulation of amnion markers AP2α and CDX2. (G) Quantification of the percentage of inner domains experessing SOX2 and CDX2 at day 4 (n=1088 structures from 3 indpendent experiments). Statistics: (D-E) Paired T-Test (G) RM two-way ANOVA with Sidak’s multiple comparisons test. *p<0.05, **p<0.01, ***p<0.001, ****p<0.0001.

To further understand potential tissue-tissue crosstalk in our human embryo model, we used the computational tool CellPhoneDB^65^ to predict ligand-receptor pairing across clusters in our single cell sequencing data (**Figure 4C, Extended** Figure 7A-B). This analysis uses the expression of curated receptor-ligand pairs across clusters to score potential tissue-tissue crosstalk. CellPhoneDB predicted that hypoblast cluster-derived BMP2/6 and extraembryonic mesenchyme-secreted BMP4 were likely mediators of tissue-tissue crosstalk, and thus potentially drivers of cell differentiation. In contrast, predicted NODAL signaling between tissues was low, further supporting the presence of a high BMP, low NODAL signaling environment in the human embryo model. When CellPhoneDB was applied to the single cell sequencing data of the three cell lines aggregated to generate the embryo model (wildtype, inducible GATA6-SOX17 and inducible GATA3-AP2γ RSeT hESCs), the inducible GATA3-AP2γ cells were predicted to be the initial source of BMP (**Extended Figure 7C**). Despite the lack of distinct trophoblast-like cells in the inducible human embryoids at later stages, aggregation of inducible GATA6-SOX17 and wildtype RSeT hESC alone (i.e. without inducible GATA3-AP2γ cells) resulted in failure to form organized structures (**Extended Figure 7D**).

To verify whether BMP signaling has an active role in the differentiation of the inner domain of the embryo model, we examined phosphorylated (p)SMAD1.5 expression. pSMAD1.5 enrichment in the OCT4-positive epiblast-like domain at days 4 and 6 post-aggregation was indicative of active BMP signaling (**Figure 4D**). This was in contrast to the low nuclear/cytoplasmic ratio of total SMAD2.3, reflecting low NODAL signaling, in the epiblast-like domain (**Figure 4E**). These findings accord with the CellPhoneDB predictions. To functionally validate the role of BMP signaling in the differentiation of the inner domain, we treated the human embryo model with the ALK1/2/3/6 (BMP receptors) inhibitor LDN193189 between 48- and 96-hours post-aggregation. LDN193189 treated structures exhibited increased maintenance of SOX2 expression in the inner domain and lesser CDX2 and AP2α upregulation at day 4 and day 6 post-aggregation, compared to untreated control structures. Supplementation with Activin-A, an agonist of SMAD2.3 signaling, resulted in a similar phenotype, though to a lesser degree (**Figure 4F-G**, **Extended Figure 7E**). These data demonstrate that the high levels of BMP activity and low levels of NODAL are key drivers of epiblast to amnion differentiation within the inner epithelial compartment of inducible human embryoids.

### Prolonged SOX17 overexpression inhibits formation of the anterior hypoblast

BMP signals are localized to the posterior of the embryo by the antagonistic action of the anterior hypoblast, which secretes inhibitors of BMP, WNT and NODAL, including CER1 and LEFTY1^2^. We have recently demonstrated that these markers of the anterior hypoblast are expressed in peri-and post-implantation human embryos cultured *in vitro*^2^. Examination of hypoblast-like cells in our inducible human embryoids revealed that neither *CER1* nor *LEFTY1* were meaningfully expressed in the HYPO/VE single cell sequencing cluster (**Figure 5A**). We therefore reanalyzed our previously published 10x single-cell RNA sequencing data from post-implantation human embryos and the previously scored transcription factor activity. This analysis revealed that SOX17 regulon activity was enriched in the *CER1*-negative hypoblast subcluster in post-implantation human embryos (**Figure 5B**)^2^. Indeed, induction of SOX17 singly or in combination with GATA6 resulted in decreased capacity to upregulate *CER1,* as compared with GATA6 overexpression alone (**Extended Figure 1C**). To test whether single induction of GATA6 changed the constitution of hypoblast-like cell subpopulations in our human embryo model, we generated hypoblast-like cells by either inducing GATA6 expression alone or in combination with SOX17. Embryoids generated with inducible GATA6 cells at day 4 post-aggregation exhibited an increased proportion of CER1-positive cells compared to embryoids generated with inducible GATA6-SOX17 cells (**Figure 5C**). By day 6 post-aggregation, CER1 expression had decreased in structures generated with GATA6-induced cells (**Extended Figure 8A**). However, the transient presence of CER1-zpositive anterior hypoblast-like cells functionally impacted the epiblast-like domain. Structures generated using hypoblast-like cells with single GATA6 induction showed a higher proportion of inner domains containing cells expressing primitive streak marker BRY/TBXT at day 6 post-aggregation as compared to structures with GATA6-SOX17 induction (**Figure 5D**). This is likely to be in response to the transient anterior hypoblast population protecting pluripotency for a longer period in these structures compared to structures generated with inducible GATA6-SOX17 cells lacking the CER1 anterior hypoblast-like domain.

**Figure 5.**
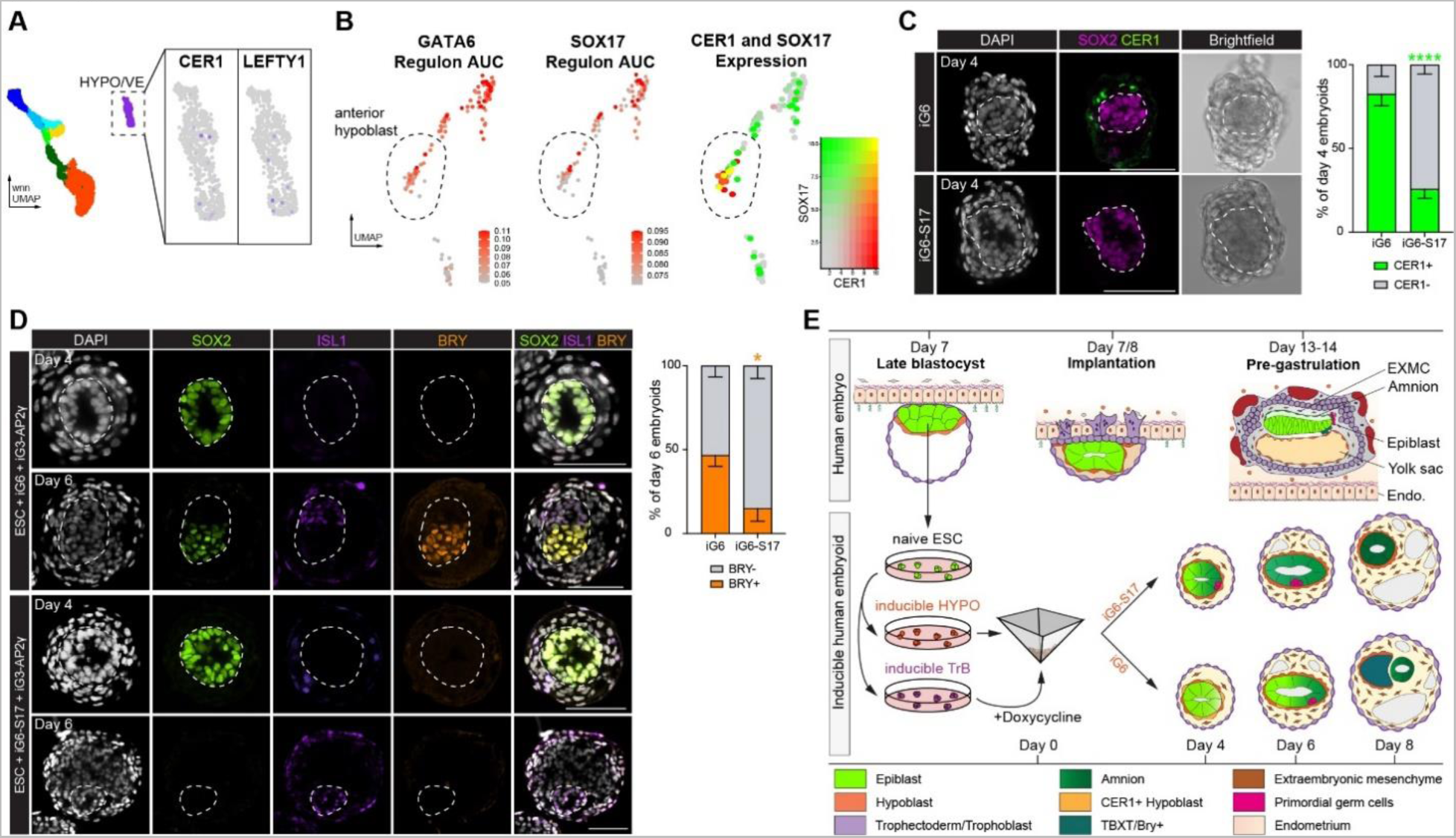
SOX17 induction is antagonistic to specification of the anterior hypoblast. (A) Expression of *CER1* and *LEFTY1* in the HYPO/VE cluster of sequenced inducible human embryoids. (B) Analysis of GATA6 and SOX17 regulon activity scored by SCENIC and *SOX17* and *CER1* co-expression in the hypoblast of post-implantation human embryos (9-11 days post-fertilization). Data from Molé et al. (C) Representative examples of inducible human embryoids and quantification showing CER1-positive cells surrounding the epiblast-like domain generated using induced GATA6 (iG6) but not induced GATA6-SOX17 (iG6-S17) dual induction of hypoblast-like cells. n=66 structures from 3 independent experiments. (D) Representative immunofluorescence staining and quantification of ISL1 and BRY/TBXT expression in inducible human embryoids generated from inducible GATA6 or inducible GATA6-SOX17 dual induction. n=29 structures from 3 independent experiments. (E) Schematic overview of inducible human embryoid formation and development compared to the human embryo. The inducible embryoids are generated by combining naïve ESCs with transgene-induced extraembryonic lineages. Embryoids self-organize and mimic aspects of human post-implantation development, including lumenogenesis, pluripotency exit, amnion and extraembryonic mesenchyme differentiation, and primordial germ cell formation. Generating embryoids with single induction of GATA6 (iG6) rather than together with SOX17 (iG6-S17) results in higher rates of anterior hypoblast differentiation, allowing for later pluripotency exit and primitive streak-like differentiation. Statistics (C-D): RM two-way ANOVA with Sidak’s multiple comparisons test. *p<0.05, **p<0.01, ***p<0.001, ****p<0.0001.

Together, these data highlight functional differences in the gene regulatory network underlying differentiation of hypoblast subpopulations of cells, where prolonged, high SOX17 expression is antagonistic to anterior hypoblast identity and instead promotes CER1-negative hypoblast identity. Thus, by modulating the hypoblast-like cells in inducible human embryoids, we reveal the capacity of the epiblast-like domain to upregulate primitive streak markers, demonstrating the value of this modular embryo model to study interactions between embryonic and extraembryonic tissues.

## Discussion

Here, we have generated a multi-lineage stem cell-derived model of the human post-implantation embryo that undergoes lumenogenesis in its epiblast-like domain and differentiation events that reflect interactions between extraembryonic-like and embryonic-like tissues. Our human embryo model forms amnion-like cells in response to BMP signaling, which progressively mature along a trajectory conserved between primates, transitioning from AP2α-positive to ISL1- and CDX2- positive to VTCN1-positive amnion cells^54, 64, 66^. These induced human embryoids specify primordial germ cell-like cells, which occurs as cells progress along the amnion differentiation trajectory, likely originating from a common AP2α-positive progenitor as reported in other *in vitro* systems^57–59^. Similarly, extraembryonic mesenchyme-like cells are specified within our embryo model, which closely resemble those of the primate embryo. The origin of extraembryonic mesenchyme in the human and non-human primate embryos remains unclear^8^. Our single cell sequencing analysis suggests a trajectory from the late epiblast-like population through a mesodermal intermediate, in line with a recently reported *in vitro* extraembryonic mesenchyme differentiation protocol^54, 62^, data from cynomolgus macaque^54^, and historical observations in rhesus macaque and human embryos^67^.

We show that modulating the transcription factor combination used to drive hypoblast-like cell induction shifts the balance of hypoblast contribution from CER1-negative hypoblast-like cells (GATA6-SOX17 induction) towards CER1-positive anterior hypoblast-like cells (GATA6 induction alone). The presence of the anterior hypoblast-like cells at day 4 post-aggregation appears to protect epiblast-like domain pluripotency for a longer period, allowing for exit from pluripotency at a developmentally later stage. This result is in agreement with the increased expression of the primitive streak marker BRY/TBXT at day 6 post-aggregation. These results suggest that RSeT hESCs acquire the capacity to differentiate into primitive streak-like cells in the embryo model over time as reported for other naïve hESC^68^. In contrast, the prolonged induction of SOX17 (together with GATA6) in hypoblast-like cells inhibits CER1 expression, leading to the earlier upregulation of pSMAD1.5 and precocious differentiation toward amnion and extraembryonic mesenchyme fates. This points to the existence of a distinct intermediate pluripotent state able to give rise to both amnion and extraembryonic mesenchyme, but not to germ layer derivatives. Thus, the human embryo model we have established here might be particularly useful to study the formation of amnion and extraembryonic mesenchyme as both lineages are not established during extended *in vitro* culture of human embryos^2, 9, 11, 13, 40, 69^.

Here, we have driven the expression of specific transcription factors to generate the two extraembryonic cell types from pluripotent hESCs and, by combining these with the parental hESC cell line, have generated a post-implantation human embryo model. Transcription factor-mediated induction of extraembryonic cells has also been used in the generation of post-implantation stem cell-derived mouse embryo models^33, 34, 46^. These methods allow for simple, robust culture of extraembryonic-like cells, which can be widely adopted. In contrast to pre-implantation blastoid models, the modular generation of embryoids from their constitutive parts (e.g. epiblast-, hypoblast- and trophoblast-like populations) will be useful in interrogating the role of specific tissues and tissue-specific gene requirements. We predict that the human embryoids we established here will become a complementary tool to address tissue-tissue crosstalk. However, the use of transcription factor overexpression to generate extraembryonic tissues may also lead to deficiencies in differentiation, as observed in models of mouse embryogenesis^33^. For example, while GATA3 and AP2γ induction drive trophoblast-like gene programs in 2D, this population is diminished in our human embryo model after aggregation and aberrantly co-expressed endodermal markers (including SOX17 and GATA6), potentially due to their peripheral position within the structure^49, 50^. Nevertheless, the GATA3-AP2γ inducible cells are required for cell aggregation, survival and organization to form the embryoids. Strikingly, induction of downstream gene regulatory networks driving both trophoblast and hypoblast lineages differed based on the initial pluripotent state. We show that this stem cell-derived model is more easily produced using peri-implantation stage, intermediate pluripotent state RSeT hESCs rather than pre-implantation stage, naïve PXGL hESCs or post-implantation stage, primed hESCs. Generating embryo-like structures from discordant pluripotent states (e.g. induction of GATA3 and/or AP2γ in naïve PXGL hESCs combined with the other two cell types from a starting RSeT condition) may lead to better recapitulation of extraembryonic fates in the future. Of note, however, PXGL hESC require extensive capacitation *in vitro* to give rise to germ layers^68^ and blastoid models generated from naïve hESCs currently fail to model post-implantation stages robustly^18–20^. Similarly, primed ESCs are described by several groups to give rise to amnion-like cells in trophoblast differentiation conditions, in contrast to naïve ESCs^29, 30, 69^. Thus, further work interrogating the epigenetic landscape and binding sites of these factors may be useful to improve strategies to generate *bona fide* extraembryonic cells.

In summary, we present a modular, tractable model of human post-implantation development that crucially includes both embryonic- and extraembryonic-like cells. This is a post-implantation model and therefore does not have the capacity to develop to form viable human embryos, as it cannot be implanted. These inducible human embryoids do not mimic stages beyond primitive streak formation, nor do they contain all cell types of the gastrulation-stage embryo^3^. However, the construction of these embryoids demonstrates a significant step towards generating integrated models of the post-implantation human embryo to allow future mechanistic studies of post-implantation development that are impossible in the real human embryo.

## Methods

### Ethics Statement

Work with human embryonic stem cells (Shef6) was carried out with approval from the UK Human Stem Cell Bank Steering Committee under approval SCSC21-38 and adheres to the regulations of the UK Code of Practice for Use of Human Stem Cell Lines. Human embryo work was regulated by the Human Fertility and Embryology Authority (HFEA) and carried out under license R0193. Ethical approval was obtained from the Human Biology Research Ethics Committee at the University of Cambridge (reference HBREC.2021.26). IVF patients at CARE Fertility, Bourn Hall Fertility and Herts & Essex Fertility Clinics were provided project-specific information at the end of their treatment. All patients were offered counselling, received no financial benefit, and could withdraw their participation at any time until the embryo had been used for research. Embryos were not cultured beyond 14 d.p.f. or the first appearance of the primitive streak.

### hESC culture

Shef6 human ESCs were cultured on Matrigel-coated plates in mTESR medium (05825, STEMCELL Technologies) at 37°C, 20% O2, 5% CO2. Plates were coated with 1.6% growth-factor reduced Matrigel (356230, BD Biosciences) dissolved in DMEM/F12 (21331-020, Life Technologies) for 1 hour at at 37°C. hESCs were passaged with TrypLE (12604013, ThermoFisher Scientific). For the first 24 hours after passaging, 10µM ROCK inhibitor Y-27632 (72304, STEMCELL Technologies) was added. Medium was changed every 24 hours. Cells were routinely tested for mycoplasma contamination by PCR (6601, Takara Bio). To convert primed hESCs to RSeT or PXGL culture conditions, cells were passaged to mitomycin-C inactivated CF-1 MEFs (3×10^3^ cells/cm^2^; GSC-6101G, Amsbio) in media consisting of DMEM/F12 with 20% Knockout Serum Replacement (10828010, ThermoFisher Scientific), 100µM β-mercaptoethanol (31350-010, Thermo Fisher Scientific), 1x GlutaMAX (35050061, Thermo Fisher Scientific), 1x non-essential amino acids, 1x penicillin-streptomycin and 10ng/ml FGF2 (Department of Biochemistry, University of Cambridge) and 10µM ROCK inhibitor Y-27632 (72304, STEMCELL Technologies). For RSeT cells, media was switched to RSeT media after 24 hours (05978, STEMCELL Technologies). Cells were maintained in RSeT and passaged as above every 4-5 days. For PXGL cells, conversion was performed as previously described^1^. Briefly, cells were cultured in 5% O2, 7% CO2 and media was switched to chemical resetting media 1 (cRM-1) consisting of N2B27 media supplemented with 1µM PD0325901 (University of Cambridge, Stem Cell Institute), 10ng/mL human recombinant LIF (300-05, PeproTech), and 1mM Valproic Acid. N2B27 contains 1:1 DMEM/F12 and Neurobasal A (10888-0222, Thermo Fisher Scientific) supplemented with 0.5x B27 (10889-038, Thermo Fisher Scientific), 0.5x N2 (made in house), 100µM β-mercaptoethanol, 1x GlutaMAX and 1x penicillin-streptomycin. cRM-1 media was changed every 48h for 4 days, after which media was changed to PXGL. PXGL media consists of N2B27 supplemented with 1µM PD0325901, 10ng/mL human recombinant LIF, 2µM Gö6983 (2285, Tocris) and 2µM XAV939 (X3004, Merck). PXGL cells were passaged every 4-6 days using TrypLE (12604013, Thermo Fisher Scientific) for 3 min. 10µM ROCK inhibitor Y-27632 and 1µL/cm^2^ Geltrex (A1413201, Thermo Fisher Scientific) were added at passage for 24 hours. For yolk sac-like cell or trophoblast differentiation, RSeT cells were passaged onto matrigel coated IBIDI chamber slides. 24 hours later, media was switched to ‘ACL’ (100ng/ml Activin-A, Qk001, QKINE, 3μM CHIR99021, 72052, STEMCELL Technologies, and 10ng/ml human LIF) for hypoblast induction or ‘PA’ (1μM PD0325901 and 1μM A83-01, 72022, STEMCELL Technologies) with or without 500nM lysophosphatidic acid – LPA (3854, Tocris).

### Generation of Inducible hESC lines

To generate Piggyback plasmids, full length coding sequences were amplified from human cell line cDNA with AttB overhangs using Phusion High-Fidelity DNA polymerase (M0530S, New England BioLabs) according to manufacturer’s instructions. Amplicons were introduced to pDONR221 entry plasmids using BP clonase (11789100, ThermoFisher Scientific), and subsequently to destination plasmids using LR clonase (11791020, ThermoFisherScientific) according to manufacturer’s instructions. hESCs were electroporated with GATA6-3XFLAG-TetOn-Zeo (entry plasmid 72922, Addgene) and/or SOX17-TetOn-Hygro or GATA3-EGFP-TetO-Hygro (Gift from Dr. Mika Drukker) and/or TFAP2C-TetOn-G418 in addition to PB-CAG-rTTA3-Bsd or PB-CAG-rTTA3-Zeo and pBase plasmid expressing PiggyBac Transposase using the Neon transfection system with the following settings: 1200V, 2 ms, 2 pulses. Two days after transfection, antibiotics were applied at a ¼ dosage and increased to final concentrations of 100µg/mL zeocin (ant-zn-1, Invitrogen), 20µg/mL blasticidin (A113903, ThermoFisher Scientific), 50µg/mL G418 (10131035, ThermoFisher Scientific) or 50µg/mL HygromycinB (10687010, ThermoFisher Scientific). Clones were generated by manually picking single colonies under a dissecting microscope. Transgene activation was triggered by addition of 1µg/mL doxycycline hyclate (D9891, Sigma). Note that AP2γ-inducible cells failed to reset in PXGL naïve conditions.

### qRT-PCR Analysis

Cell pellets were harvested and RNA was extracted using the Qiagen RNeasy kit following manufacturer’s instructions. Reverse Transcriptase reaction was performed with 1µg RNA with random primers (C1181, Promega), dNTPs (N0447S, New England BioLabs), RNAse inhibitor (M0314L, New England Biolabs, and M-MuLV reverse transcriptase (M0253L, New England Biolabs). RT-qPCR was performed using Power SYBR Green PCR Master Mix (4368708, ThermoFisher Scientific) on a Step One Plus Real-Time PCR machine (Applied Biosystems). The following program was used: 10 minutes at 95°C followed by 40 cycles of 15s at 95°C and 1 minute at 60°C. Single melt curves were observed for all primers used in this study.

### Generation of Inducible Human Embryoids

To generate the three-dimensional stem cell-derived model of the post-implantation embryo, RSeT cells between 2 and 6 passages post-conversion to RSeT media were passaged as normal; the following day (Day -3) the media for extraembryonic-like cells (induced GATA6, induced GATA6-SOX17 or induced GATA3-AP2γ) was changed to N2B27 with 5% Knockout Serum Replacement and 1µg/mL DOX. This media was refreshed every 24 hours for 3 days. On day 0 (the day of aggregation), an Aggrewell dish (34415, STEMCELL Technologies) was prepared by pre-coating with anti-adherance solution (07010, STEMCELL Technologies) and centrifuging at 2000g for 5 minutes. Wells were washed twice with PBS prior to the addition of experiment media. This media consists of N2B27 with 5% knockout serum replacement, 1µg/mL doxycycline and 10µM Y-27632. Induced cells and wild type ESCs were enzymatically dissociated 1 hour after addition of 10µM Y-27632 to wells containing cells for inducible human embryoid generation. Dissociated cells were pelleted and resuspended in experiment media and placed in gelatin-coated wells for MEF depletion. After 15-30 minutes cells were counted, mixed, and plated into an Aggrewell dish with a final calculation of 8 wildtype-ESC, 8 hypoblast-like and 16 trophoblast-like cells plated per microwell in the Aggrewell. On day 1, the media was subjected to two two-third changes of N2B27 with 5% knockout serum replacement and 1µg/mL doxycycline. On day 2, Aggrewells were subjected to a half change with hIVC1 media with 1.25 µg/mL hIGF1 (78022.1 STEMCELL Technologies) and 1µg/mL doxycycline. hIVC1 media consists of Advanced DMEM/F12 (12634-010 ThermoFisher Scientific) supplemented with 20% inactivated FBS (10270106, ThermoFisher Scientific), 1x Glutamax, 1X NEAA, 1x Essential AA, 1x ITS-X, 25U/mL Pen/Strep, 1.8mM Glucose (G8644, Sigma-Aldrich), 0.22% sodium lactate (L7900, Sigma-Aldrich), 8nM β-estradiol (50-28-2, Tocris) and 200ng/mL progesterone (P0130, Sigma-Aldrich). This media is used in half changes each day from day 3. On day 4, aggregates were manually picked using a mouth pipette under a dissecting microscope into individual wells of ultra-low attachment 96 well plates (CLS7007, Corning) in hIVC1 media with IGF1 and doxycycline as above for subsequent culture.

### Immunostaining and image analysis

Samples were washed with phosphate-buffered saline (PBS) and fixed in 4% Paraformaldehyde (PFA; 1710, Electron Microscopy Sciences) at room temperature for 20 minutes. Samples were washed 3 times with PBS + 0.1% (vol/vol) Tween-20 (PBST) and incubated with 0.3% (vol/vol) Triton X-100 (T8787, Sigma Aldrich) + 0.1mM glycine (BP381-1, Thermofisher Scientific) in PBS at room temperature for 30 minutes. Samples were blocked in blocking buffer (PBST with 5% (w/vol) BSA, A9418, Sigma), then incubated with primary antibodies diluted in blocking buffer overnight at 4°C. See table 1 for a list of primary antibodies. Samples were washed three times in PBST and incubated with fluorescently conjugated AlexaFlour secondary antibodies (Thermofisher Scientific, 1:500) and DAPI (D3571, ThermoFisher Scientific, 1μg/ml) diluted in blocking buffer for 2 hours at room temperature. For pSMAD1.5 quantification, OCT4-positive or GATA6-positive nuclei were isolated and the fluorescent intensity of pSMAD1.5 was quantified. For SMAD2.3 quantification, OCT4-positive or GATA6-positive (excluding the outermost GFP+ cell layer) nuclei fluorescence intensity was quantified, as well as cytoplasmic fluorescence intensity. Data is presented as the ratio of nuclear:cytoplasmic fluorescence intensity. Immunofluorescence images were analyzed using FIJI.

**Table 1:** List of primary antibodies used in this study.

| Target | Species | Company | Catalogue # | RRID |
| --- | --- | --- | --- | --- |
| AP2-alpha | Mouse | Santa Cruz Biotechnology | sc-12726 | AB_667767 |
| AP2-gamma | Goat | R&D Systems | AF5059 | AB_2255891 |
| Brachyury | Goat | R&D Systems | AF2085 | AB_2200235 |
| CDX2 | Mouse | BioGenex | MU392-UC | AB_2335627 |
| Cerberus 1 | Goat | R&D Systems | AF1075 | AB_2077228 |
| Cytokeratin 7 | Mouse | Agilent | M7018 | AB_2134589 |
| E-Cadherin | Mouse | BD Biosciences | 610182 | AB_39581 |
| EOMES | Rabbit | Abcam | ab23345 | AB_778267 |
| FOXA2 | Goat | R&D Systems | AF2400 | AB_2294104 |
| FOXF1 | Rabbit | ThermoFisher Scientific | PA5-50516 | AB_2607993 |
| GATA3 | Goat | Abcam | ab199428 | AB_2819013 |
| GATA6 | Goat | R&D Systems | AF1700 | AB_2108901 |
| GATA6 | Rabbit | Cell Signaling Technology | 5851 | AB_10705521 |
| GFP | Chicken | Abcam | ab13970 | AB_300798 |
| GFP | Rat | Nacalai USA | GF090R | AB_2314545 |
| HAND1 | Mouse | DSHB | PCRP-HAND1-2A9 | AB_2618668 |
| ISL1 | Mouse | DSHB | PCRP-ISL1-1A9 | AB_2618775 |
| N-Cadherin | Mouse | Abcam | ab98952 | AB_10696943 |
| NANOG | Rabbit | Cell Signaling Technology | 4903 | AB_10559205 |
| OCT3/4 | Mouse | Santa Cruz Biotechnology | sc-5279 | AB_628051 |
| OTX2 | Goat | R&D Systems | AF1979 | AB_2157172 |
| phosp-Smad1/5 | Rabbit | Cell Signaling Technology | 9516S | AB_491015 |
| Smad2/3 | Rabbit | Cell Signaling Technology | 8685S | AB_10889933 |
| SOX17 | Goat | R&D Systems | AF1924 | AB_355060 |
| SOX2 | Rat | ThermoFisher Scientific | 14-9811-82 | AB_11219471 |
| TBX20 | Mouse | R&D Systems | MAB8124 | AB_2255728 |
| VTCN1 | Rabbit | Abcam | ab209242 | AB_2801513 |

**Table 2:**
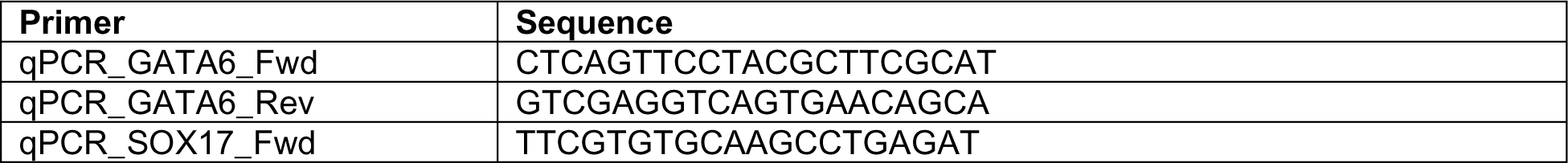
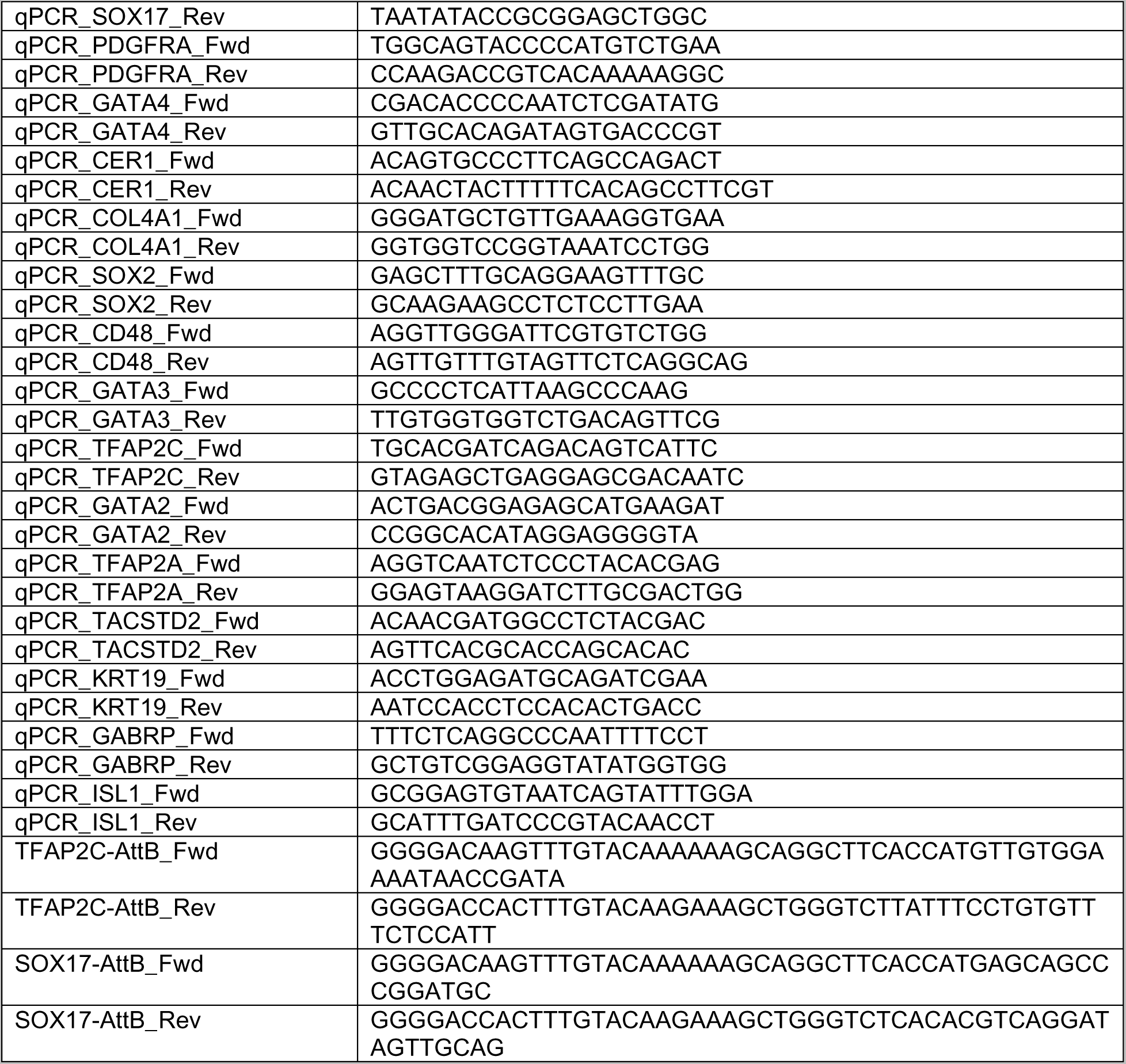
List of primers used in this study.

### Human Embryo Thawing and Culture

Human embryos were thawed and cultured as described previously^2, 3^. Briefly, cryopreserved human blastocysts (5 or 6 d.p.f.) were thawed using the Kitazato thaw kit (VT8202-2, Hunter Scientific) according to the manufacturer’s instructions. The day prior to thawing, TS solution was placed at 37°C overnight. The next day, IVF straws were submerged in 1mL pre-warmed TS for 1 min. Embryos were then transferred to DS for 3 min, WS1 for 5 min, and WS2 for 1 min. These steps were performed in reproplates (REPROPLATE, Hunter Scientific) using a STRIPPER micropipette (Origio). Embryos were incubated at 37°C and 5% CO2 in normoxia and in pre-equilibrated human IVC1 supplemented with 50ng/mL Insulin Growth Factor-1 (IGF1) (78078, STEMCELL Technologies) under mineral oil for 1-4h to allow for recovery. Following thaw, blastocysts were briefly treated with acidic Tyrode’s solution (T1788, Sigma) to remove the zona pellucida and placed in pre-equilibrated human IVC1 in 8 well µ-slide tissue culture plates (80826, Ibidi) in approximately 400 µL volume per embryo per well. Half media changes were done every 24 hours.

### Statistics

Statistical analyses were performed using Graphpad Prism 9. Sample sizes were not predetermined, and the authors were not blinded to conditions. All experiments were performed independently at least twice. Within plots, all data is presented as mean ± SEM.

### Collection, Generation and Sequencing of single-nuclei ATAC/RNA 10x Libraries

To collect post-implantation embryo models for single-cell sequencing, correctly organized embryoids at day 4, 6 and 8 were picked visually and washed through PBS twice in a 4-well dish before transfer to TrypLE. Samples were agitated by pipetting every 5 minutes for 10-20 minutes until dissociated. Enzymatic activity was inactivated through addition of 20% FBS in PBS at 2x volume. Cells were collected in a falcon tube, pelleted, and resuspended in freeze buffer consisting of 50 mM Tris at pH 8.0 (15-567-027, Fisher Scientific), 25% glycerol (G5516, Sigma-Aldrich), 5 mM Mg(OAc)2 (63052, Sigma-Aldrich), 0.1mM EDTA (15575020, ThermoFisher Scientific), 5mM DTT (R0861, ThermoFisher Scientific), 1x protease inhibitor cocktail (P8340, Sigma-Aldrich), 1:2500 dilution of superasin (AM2694, Invitrogen). For cell lines, 10,000 cells were counted, pelleted, and resuspended in the freeze media above before slowfreezing at -80°C.

For nuclei isolation and library construction, low input nuclei isolation protocol from 10x Genomics was performed. Briefly, frozen cell pellets were thawed in a 37°C water bath for 30 seconds, centrifuged (500g for 5 minutes at 4°C) to pellet the cells then the supernatant were aspirated. The cell pellets were washed with 200 µl 1x PBS with 0.04% BSA twice, centrifuged, and supernatant was aspirated between washes. Subsequently, chilled lysis buffer (45 µl per sample) was added to the washed cell pellet and placed on ice for 3 minutes, then wash buffer (50 µl per sample) was added. Washed isolated nuclei were resuspended in a diluted nuclei buffer. In this study, the isolated nuclei were resuspended in 5 µl of diluted nuclei buffer and were directly added to the transposition reaction. In all following steps, 10x Genomics’ Single Cell Multiome ATAC and Gene expression protocol were followed according to manufacturer’s specifications and guidelines. The final libraries were loaded on the NextSeq 2000 using P2 100 cycle kit at 650 pM loading concentration with paired end sequencing following the recommended sequencing reads from 10x Genomics (28/10/10/90 cycles for gene expression libraries and 50/8/24/49 cycles for ATAC libraries).

### Single Cell Sequencing Analysis

#### Processing and Quality Control

Raw reads were analyzed using the CellRangerARC pipeline to generate ATAC and RNA fastq files for each sample, and then to align genomic and transcriptomic reads. Matrices were then read into Seurat^4^ and Signac^5^ using the Read10X_h5 command. For ATAC-seq data, peaks from standard chromosomes were used and peaks were additionally called using macs2 to add an additional Signac assay. Cells with >500 RNA UMI counts, <20% mitochondrial reads, >500 ATAC reads, TSS enrichment >1 and were called as singlets using scDblFinder^6^ were retained for downstream analysis. For UMAP projections, SCTransform was used for RNA counts with percent mitochondrial counts and cell cycle scores regressed. PCA and LSI graphs were used to generate a weighted nearest neighbor (wnn) embedding which accounts for both modalities. chromVAR^7^ was run to calculate motif accessibility score on the peaks assay.

#### Comparisons to Published Datasets

scmap^8^ was used to project cell labels from other single cell datasets onto post-implantation embryo model transcriptional data. All reference data used was publicly available with published cell type annotations. Cynomolgus monkey gene names were converted to hgnc gene symbols using biomaRt. For data generated with smart-seq2 or other non-UMI-based single cell sequencing methods, the scmapCluster method was used with a similarity threshold of 0.5. For UMI-based methods the scmapCell followed by scmapCell2Cluster method was used with w=2. Upon cell type assignment and processing for the cell lines sequenced, a logistic regression framework was applied to project cell line data onto published single cell data and to project published cluster annotations onto post-implantation embryo model clusters, resulting in a quantitative measure of predicted similarities^9^. Here, only differentially expressed genes (produced using Seurat’s FindAllMarkers function on course cell assignments (which collapsed amnion and mesodermal clusters) were used.

#### Multivelo RNA/Chromatin Velocity

We applied a recently published method for velocity calculations that accounts for both single cell ATAC and RNA data^10^. Multivelo was run on all cells which passed the QC and processing described above. Analysis was based on available vignettes with 1000 highly variable genes and the ‘grid’ method.

#### CellPhoneDB Analysis

CellPhoneDB 2.0 was used with default settings to assess potential tissue signaling crosstalk^11^. Course cell assignments which collapsed separated amnion (AM-1, AM-2, AM-3) and mesoderm (MESO-1, MESO-2) clusters was used for simplicity. Selected significant interactions were plotted as dot plots.

#### Reanalysis of human *in vitro* cultured embryo datasets

Previously published data from Yan et al., Blakely et al., Petropolous et al., Zhou et al., and Xiang et al., were realigned to the hg38 human genome using kallisto or kb-bustools^12, 13^. Datasets which were not sequenced with UMI-based technologies were normalized using quminorm to quasi-umis^14^. Using SCTransform-based integration, datasets were combined to generate a single-cell RNA-seq dataset of human embryos spanning zygote to day 14 post-fertilization^2, 15–19^. Cells were clustered and identities assigned based on previous annotations and canonical marker expression. SCENIC was used with default settings in R, and the AUC-regulon table used to generate a new assay in the Seurat object. Using this assay, the epiblast, hypoblast and trophoblast lineages were then compared using Seurat’s FindMarkers function to identify predicted differentially active regulons. These factors were then plotted in relation to each other in Cytoscape.

#### Data Availability Statement

Data generated in this study is available under Gene Expression Omnibus accession code GSE218314. For the purpose of revew this dataset can be accessed with the token ijkhciquzhubbqt.

Previously published data is publicly available: Human data

Molè et al., 2021: ArrayExpress E-MTAB-8060

Xiang et al., 2020: Gene Expression Omnibus GSE136447

Zhou et al., 2019: Gene Expression Omnibus GSE109555

Petropoulos et al., 2016: ArrayExpress E-MTAB-3929

Blakely et al., 2015: Gene Expression Omnibus GSE66507

Yan et al., 2013: Gene Expression Omnibus GSE36552

Cynomolgus Monkey

Yang et al., 2021: Gene Expression Omnibus GSE148683

Ma et al., 2019: Gene Expression Omnibus GSE130114

Nakamura et al., 2016: Gene Expression Omnibus GSE74767

#### Code Availability Statement

Code to used to analyze these data is available at https://github.com/bweatherbee/human_model (to be made public upon publication)

## Acknowledgements

The authors are grateful to CARE Fertility, Herts and Essex Fertility Centre, Bourn Hall Fertility Clinic and King’s Fertility Clinic for their collaboration in donation of supernumerary human embryos. The authors thank all members of the M.Z-G and Boroviak labs, Gianluca Amadei for thoughtful comments and Marta Shahbazi and David Glover for feedback on the manuscript. This work is supported by Wellcome Trust, Open Atlas and NOMIS award grants to M.Z-G, Allen Discovery Center for Cell Lineage Tracing grants to J.S. and M.Z-G., in addition to individual funding from the Gates Cambridge Trust (to B.A.T.W.) and Leverhulme Trust Early Career Fellowship (to C.W.G.). J.S. is an investigator of the Howard Hughes Medical Institute.

## Author Contributions

B.A.T.W. and C.W.G. designed, carried out, and analyzed experiments. B.A.T.W., C.W.G. and L.K.I-S. performed human embryo work. N.H. and R.M.D. prepared 10x multiome libraries and performed RNA and ATAC sequencing supervised by J.S.. B.A.T.W. analyzed single-cell sequencing data. C.W.G. and B.A.T.W. wrote the manuscript with input from all co-authors. M.Z-G conceived and supervised the project.

## Extended Data

**Extended Figure 1 (related to Fig 1).**
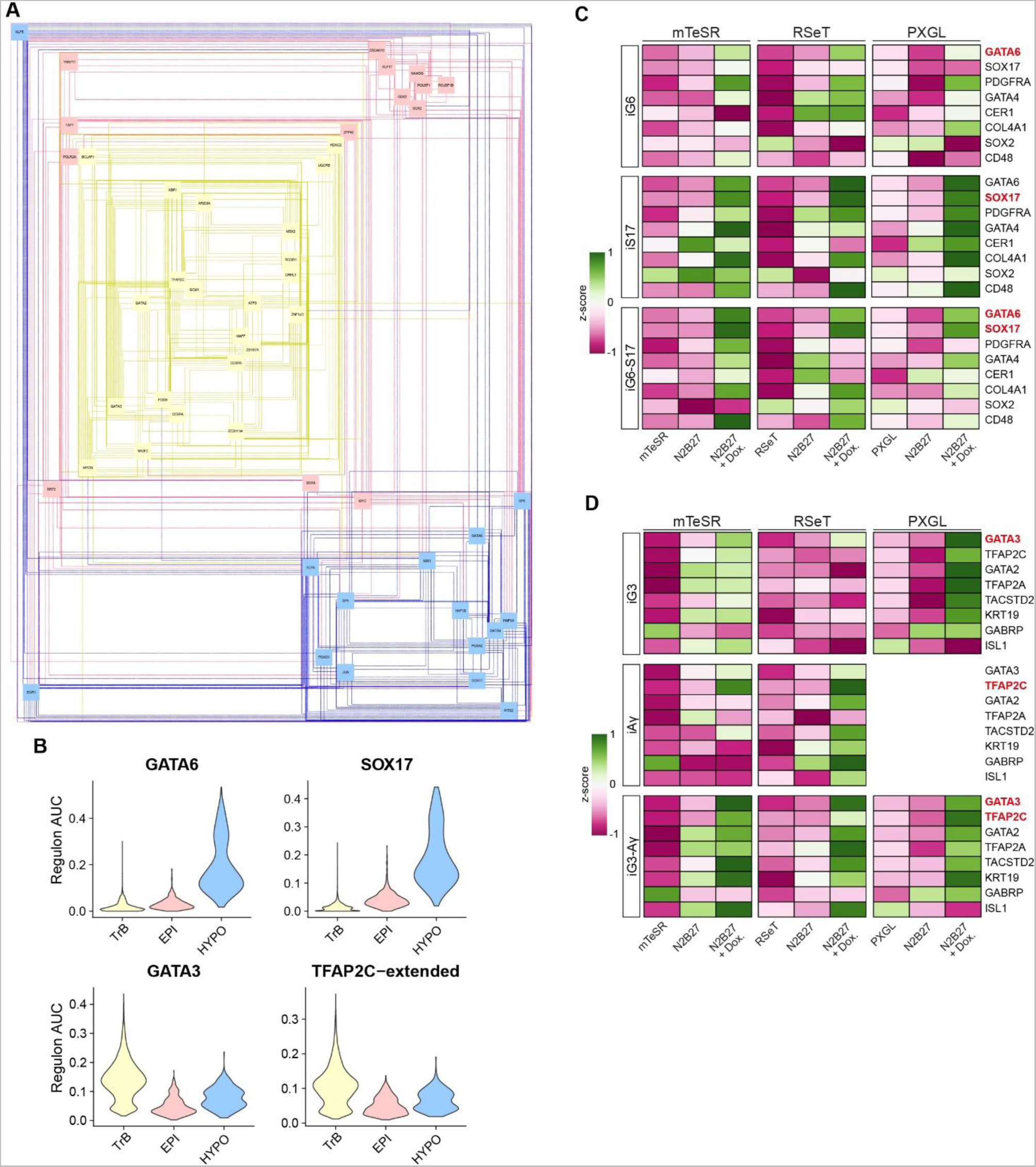
Sequencing analysis and gene regulatory network generation to guide transgene selection. (A) Inferred epiblast, hypoblast and trophoblast gene regulatory network generated by SCENIC during peri-implantation human embryo development. See methods for details on datasets and processing. (B) Regulon activity scored by SCENIC for hypoblast markers GATA6, SOX17 and TrB markers GATA3 and TFAP2C. (C) qRT-PCR analysis after 3 days of doxycycline-induction of induced GATA6 (iG6), induced SOX17 (iS17) or induced GATA6-SOX17 (iG6-S17) singly and together from three pluripotent states. N=3 technical replicates from 3 biological replicates. (D) qRT-PCR analysis after 3 days of DOX-induction of induced GATA3 (iG3), induced AP2γ (iAγ) or induced GATA3-AP2γ (iG3-Aγ) from multiple pluripotent starting states. N=3 technical replicates for 3 biological replicates. Note, for (C) and (D) the induced transgenes are marked in red.

**Extended Figure 2 (related to Fig 1).**
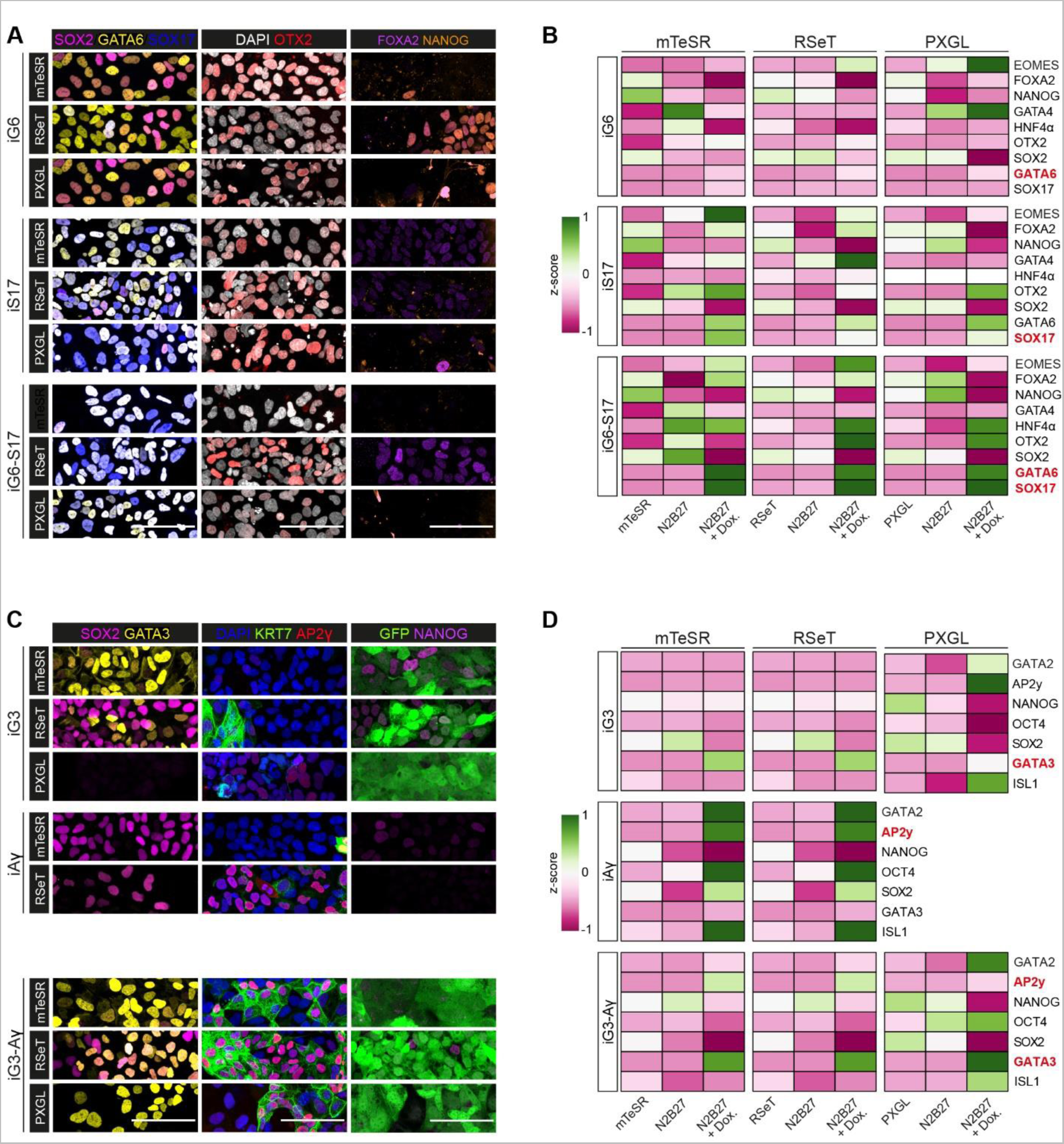
Immunofluorescence analysis of cardinal marker genes of hypoblast and trophoblast after doxycycline induction across pluripotent states. (A) Immunofluorescence analysis of inducible GATA6 (iG6), inducible SOX17 (S17) or inducible GATA6-SOX17 (iG6-S17) cells after 3 days induction from multiple pluripotent states. (B) Quantification of A. (C) Immunofluorescence analysis of inducible GATA3 (iG3), inducible AP2γ (iAγ) or inducible GATA3-AP2γ (iG3-Aγ) after 3 days induction from multiple pluripotent states. Note, for (B) and (D) the induced transgenes are marked in red. For all, N=3 technical replicates from 2 independent experiments.

**Extended Figure 3 (related to Fig 1).**
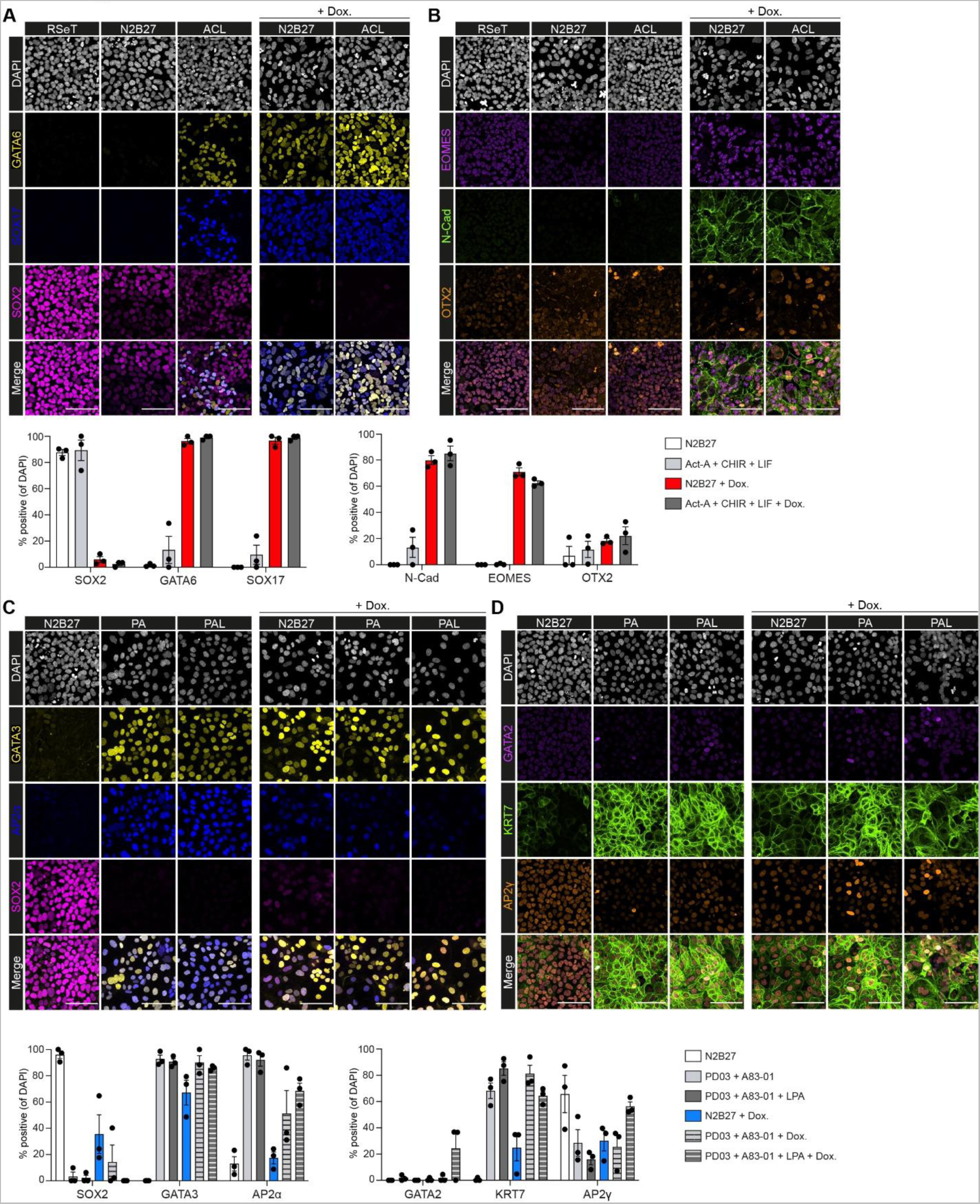
Comparison of TF-mediated induction with published directed differentiation methods. (A) Comparison and quantification of GATA6, SOX17 and SOX2 after yolk sac-like cell (Activin-A, CHIR99021 and LIF) directed differentiation, doxycycline-mediated induction in inducible GATA6-SOX17 cells or both. Cells were differentiated from RSeT conditions. (B) Comparison and quantification of EOMES, N-Cadherin and OTX2 after yolk-sac like cell (Activin-A, CHIR99021 and LIF) directed differentiation, doxycycline-mediated induction in inducible GATA6-SOX17 cells or both. N=2994 cells. (C) Comparison and quantification of GATA3, AP2α and SOX2 after PA (PD0325901 and A83-01) or PAL (PD0325901, A83-01 and LPA) directed differentiation, doxycycline-mediated induction in inducible GATA3-AP2γ cells or both. (D) Comparison and quantification of GATA2, KRT7 and AP2γ after PA (PD0325901 and A83-01) or PAL (PD0325901, A83-01 and LPA) directed differentiation, doxycycline-mediated induction in inducible GATA3-AP2γ RSeT cells or both. N=2368 cells. Differentiation was carried out on hESC in RSeT conditions.

**Extended Figure 4 (related to Fig 1).**
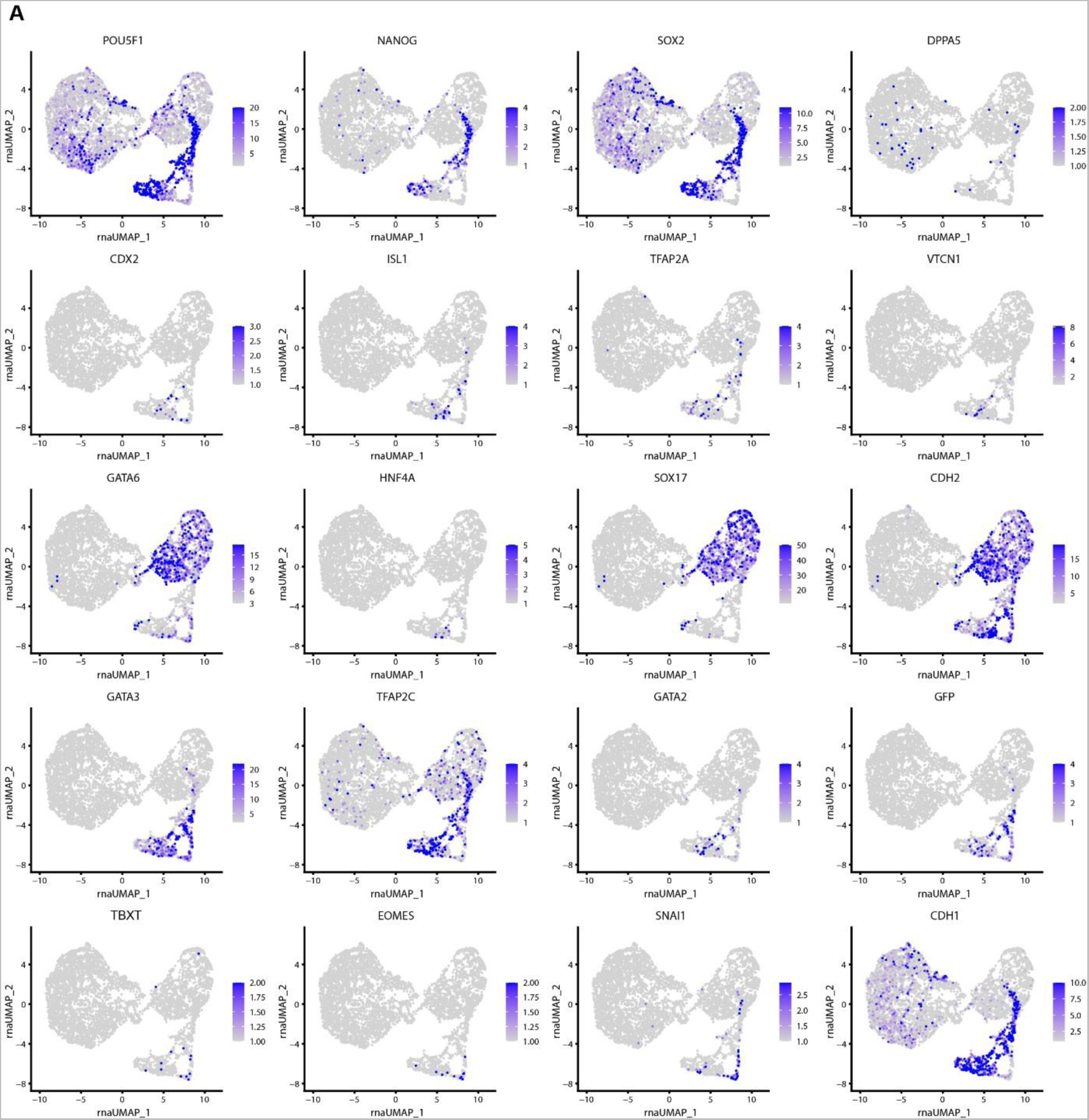
Expression of cardinal marker genes after three days of induction from RSeT cells. (A) UMAP projections of selected genes after 3 days of doxycycline-induction from sequencing of wildtype, inducible GATA6-SOX17 and inducible GATA3-AP2γ RSeT hESC populations. Induction was carried out from hESC in RSeT conditions.

**Extended Figure 5 (related to Fig 3).**
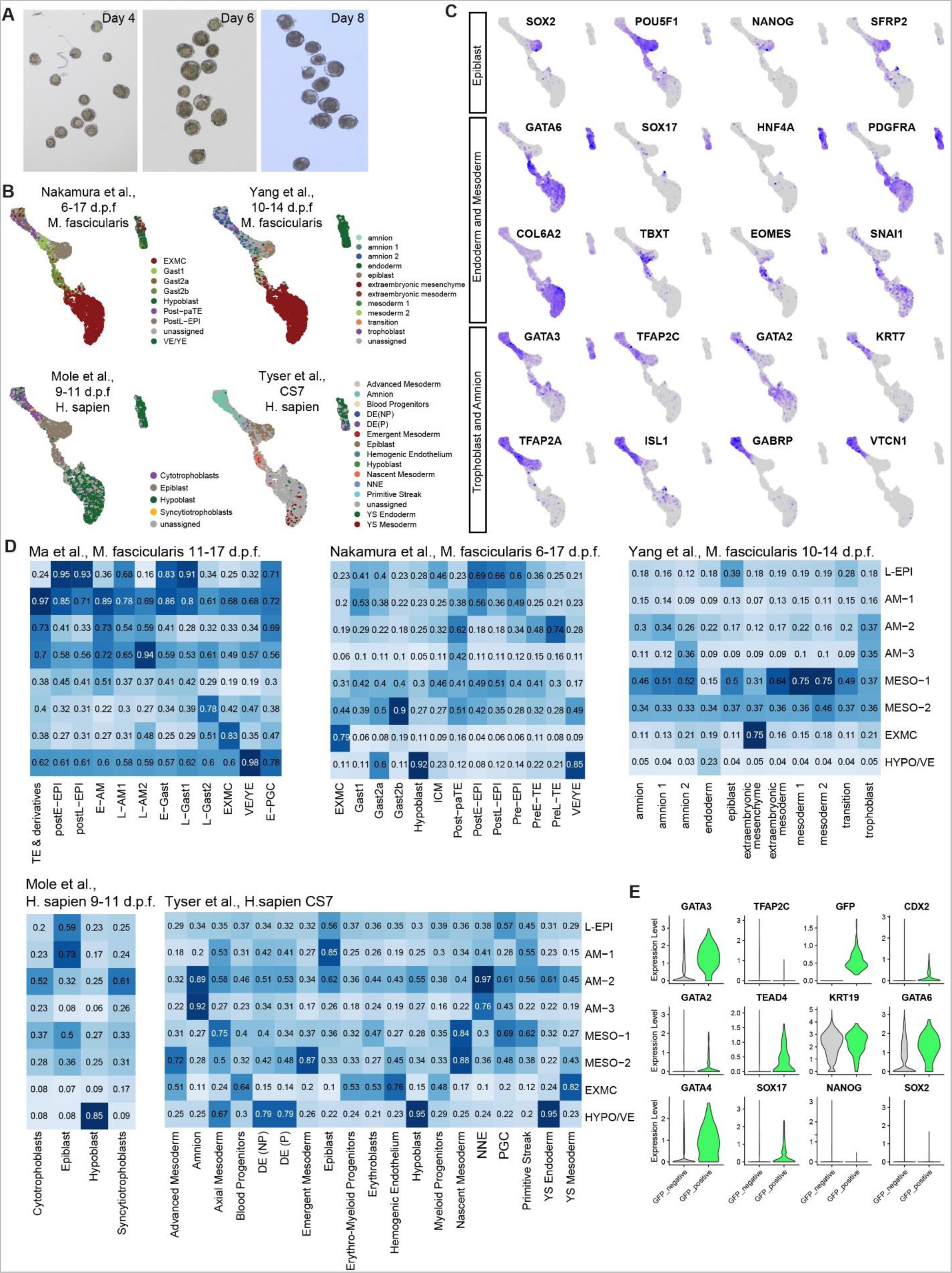
Post-implantation human model cluster identification and comparison to human and cynomolgus monkey datasets. (A) Brightfield image of inducible human embryoids selected at days 4, 6 and 8 for sequencing (n=12 at each stage). Note the presence of an inner domain surrounded by two concentric domains. (B) Cardinal marker gene expression for Epiblast, Endoderm and Mesoderm, and Trophoblast and Amnion within the stem cell derived model. (C) scmap projection of inducible human embryoid cells onto cynomolgus macaque (*M. fasicularis*) and human datasets (*H. sapien*) spanning peri-implantation to gastrulation stages. (D) Logistic regression analysis projecting annotated clusters from cynomolgus macaque (*M. fasicularis*) and human datasets (*H. sapien*) spanning peri-implantation to gastrulation stages onto post-implantation human embryo model clusters. Cynomolgus data from Ma et al., 2019, Nakamura et al., 2016, Yang et al., 2021; Human data from Molè et al., 2021 and Tyser et al., 2021. (E) Violin plots of gene expression in GFP-negative versus GFP-positive cells derived from induced GATA3-AP2γ cells from inducible human embryoid sequencing dataset.

**Extended Figure 6 (related to Fig 3).**
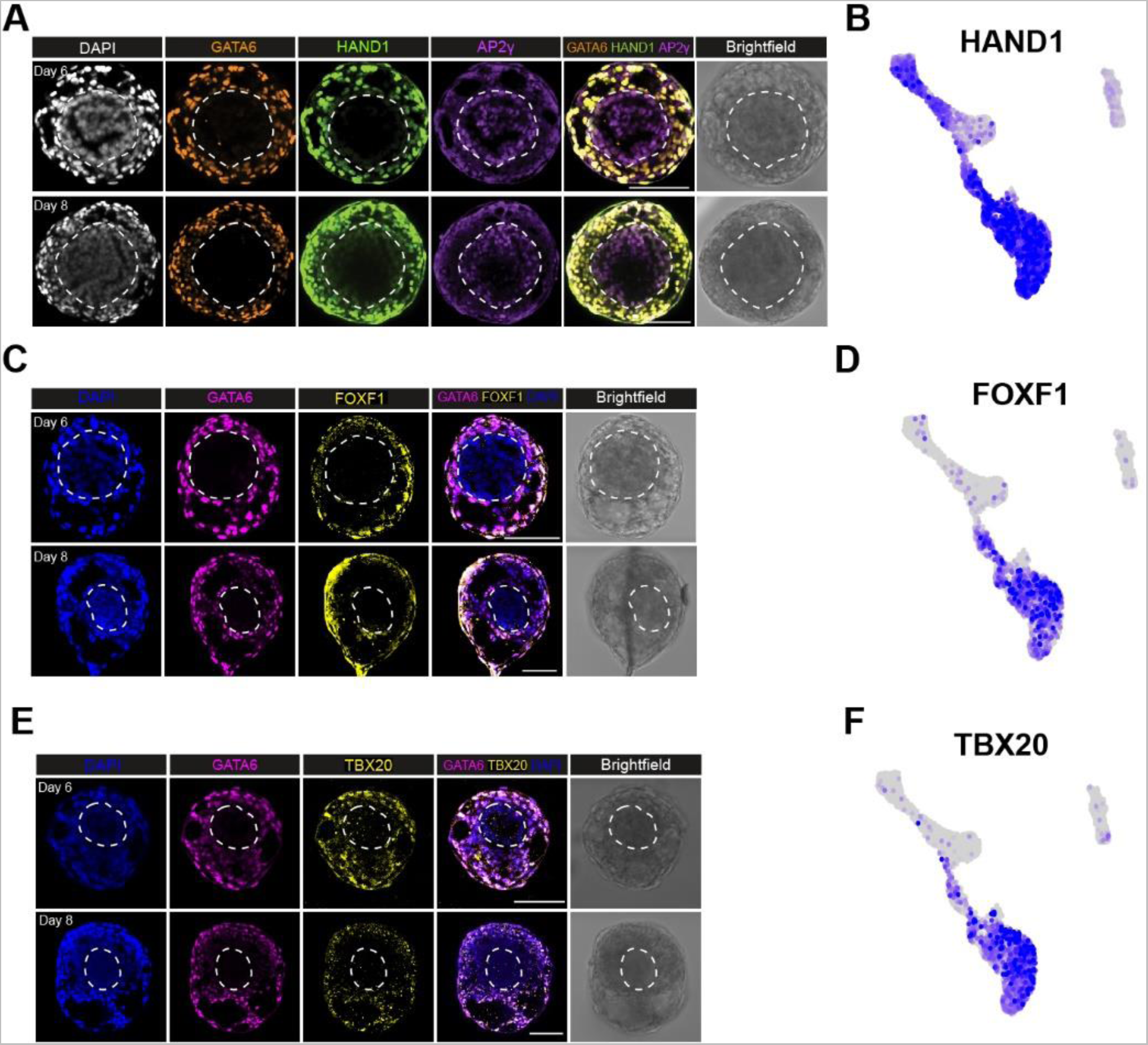
Upregulation of HAND1 in extraembryonic mesenchyme and amnion trajectories. (A) Immunofluorescence of HAND1 demonstrating expression in GATA6-positive cells (putative extraembryonic mesenchyme) and upregulation between days 4 and 6 in putative amnion (AP2γ-positive). (B) Expression of *HAND1* in the inducible human embryoid single cell sequencing dataset. (C) Immunofluorescence of FOXF1 demonstrating high expression in a subset of GATA6-positive cells (putative extraembryonic mesenchyme). (D) Expression of *FOXF1* in the inducible human embryoid single cell sequencing dataset. (E) Immunofluorescence of TBX20 demonstrating high expression in a subset of GATA6-positive cells (putative extraembryonic mesenchyme). (F) Expression of *TBX20* in the inducible human embryoid single cell sequencing dataset.

**Extended Figure 7 (related to Fig 4).**
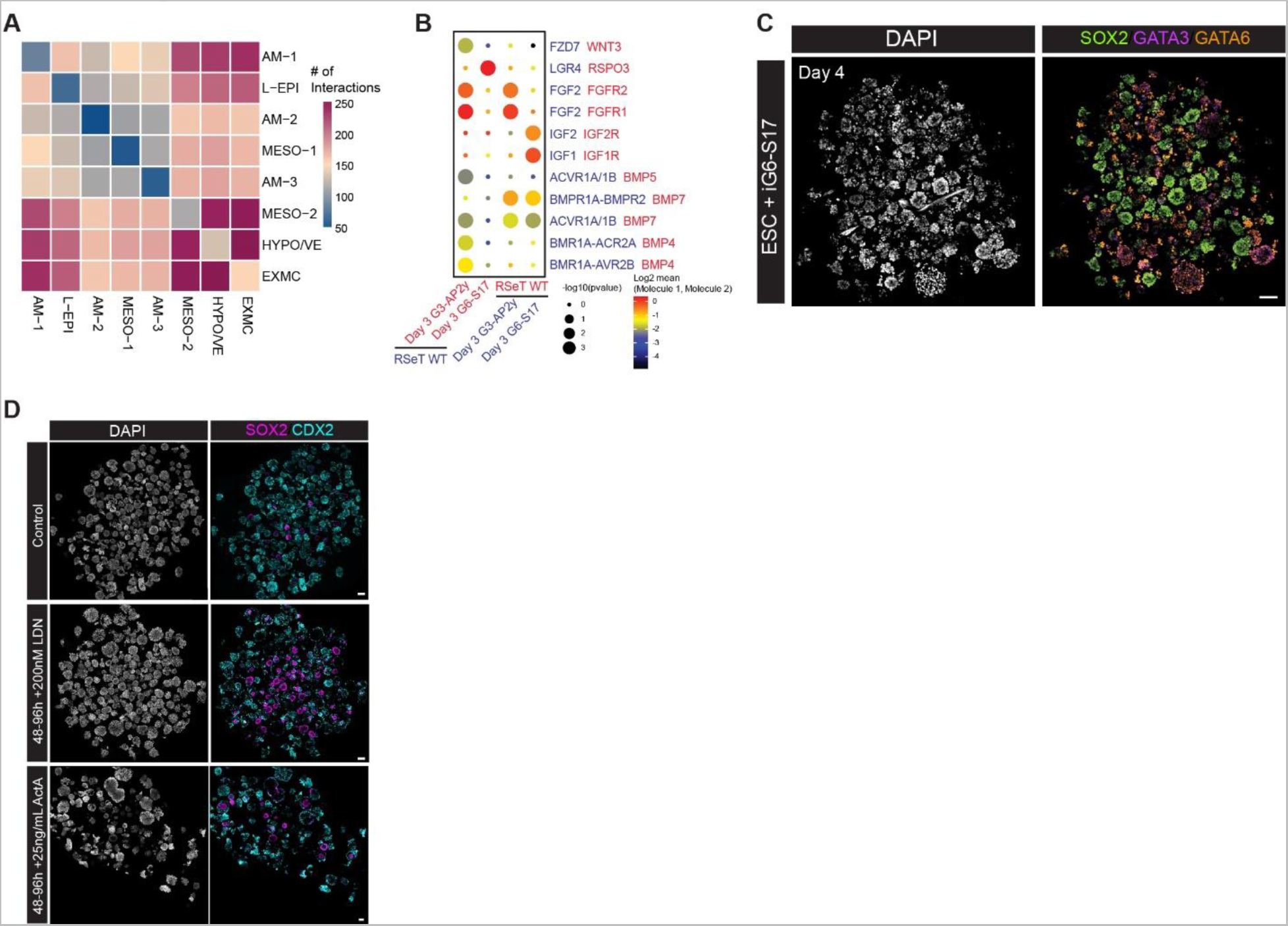
Importance of BMP and inducible GATA3-AP2γ cells in generating inducible human embryoids. (A) Predicted interaction map between clusters generated with CellPhoneDB. (B) Predicted interactions of inducible GATA6-SOX17 (G6-S17) and inducible GATA3-AP2γ (G3-AP2γ) cells after 3 days induction with wildtype RSeT hESCs, which are the cell types aggregated to generate inducible human embryoids. (C) Inducible human embryoids do not form if induced GATA3-AP2γ cells are excluded. (D) Overview of whole Aggrewells demonstrating the effect of BMP inhibition of NODAL activation. Note the significant increase in well-organized structures expressing SOX2 after BMP inhibition.

**Extended Figure 8 (related to Fig 5).**
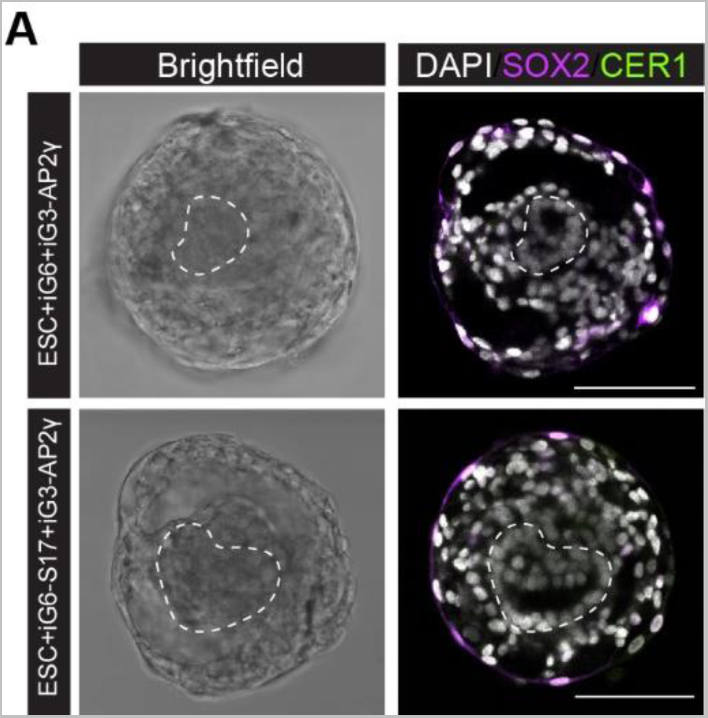
CER1 expression is downregulated upon extend culture of inducible human embryoids. (A) Immunofluorescence of CER1 and SOX2 at day 6 post-aggregation demonstrates downregulation of both SOX2 and CER1 at this stage in structures generated with both inducible GATA6 (iG6) or inducible GATA6-SOX17 (iG6-S17) hypoblast-like cells together with wildtype ESCs and inducible GATA3-AP2γ (iG3-AP2γ cells.

**Extended Table 1: Differentially Expressed Transcripts, Differentially Accessible Motifs, and Differentially Accessible Peaks between Cell Lines**

**Extended Table 2: Differentially Expressed Transcripts, Differentially Accessible Motifs, and Differentially Accessible Peaks between inducible Human Embryoid Course Cell Assignments**

